# Distinct atypical chemokine receptor 1 determinants underlie bacterial toxins recognition and pore formation

**DOI:** 10.64898/2026.09.07.749789

**Authors:** Aurélien Fouillen, Elodie Logerot, Zainab Ammar, Tristan Girbau, Isabelle Brabet, Omolade O. Otun, Rym Ben Boubaker, Camille Le Drezen, Maria Bacia-Verloop, Ambroise Desfosses, Leyrat Cédric, Sébastien Granier, Cherine Bechara

## Abstract

Atypical chemokine receptor 1 (ACKR1) is one of the most promiscuous receptors in the human chemokine system, engaging structurally diverse chemokines through a conformationally flexible N-terminal tail. This same interface is exploited by pathogens, including *Plasmodium vivax* and *Staphylococcus aureus* (SA), via a compact sulfotyrosine code. Among pathogenic proteins recognizing ACKR1, the SA leukocidin pair HlgAB is a notable exception. HlgAB-mediated pore formation is only weakly competed by chemokines, the Duffy binding protein, or antibodies targeting the receptor’s N-terminus, leaving open how HlgAB engage ACKR1. Combining structural biology approaches with cell-based assays, we show that HlgA and HlgB engage ACKR1’s sulfated N-terminus with distinct affinities and site hierarchies, with a single higher-affinity site for sulfated tyrosine 41 present in HlgA but absent in HlgB. Unexpectedly, this N-terminal engagement is dispensable for pore formation; productive lysis instead requires a separate interaction between the toxins and ACKR1’s extracellular vestibule. These results define a two-step recognition mechanism, toxin capture by the sulfotyrosine N-terminus followed by vestibule-dependent pore formation, and extend the view that ACKR1’s promiscuity arises from distributed, ligand-specific use of multiple receptor surfaces rather than a single adaptable interface.

## Introduction

Atypical chemokine receptor 1 (ACKR1), also known as the Duffy antigen receptor for chemokines (DARC), is one of the most promiscuous receptors in the human chemokine system, binding the majority of CC and CXC chemokines while remaining uncoupled from canonical G protein signaling^1–4^. This ligand promiscuity underlies its physiological role in regulating chemokine gradients that direct leukocyte trafficking, and is also the basis for its exploitation by pathogens^5^. ACKR1 is the obligate invasion receptor for the malaria parasites *Plasmodium vivax* and *P. knowlesi*^6–9^, and it is hijacked by *Staphylococcus aureus*, which secretes the bicomponent pore-forming leukocidins LukED and HlgAB to lyse red blood cells and endothelial cells as part of its strategy for iron acquisition and systemic dissemination^10–12^. Since unrelated pathogens have converged on ACKR1, the very receptor that regulates host chemokine biology, defining the structural basis for how this single receptor accommodates such diverse ligands is essential to understanding both chemokine biology and infectious diseases.

Structural insight into how ACKR1 achieves this remarkable ligand versatility has advanced considerably in recent years, but a coherent picture has yet to emerge. Cryo-electron microscopy (cryo-EM) structures of ACKR1 bound to the CC chemokine CCL7 and, more recently, the CXC chemokine CXCL8 show that both ligands engage the receptor almost exclusively through its N-terminal tail, docking superficially against a partially formed, non-canonical pocket rather than inserting into the transmembrane bundle as seen for prototypical chemokine receptors^13,14^. This single-site, N-terminus-dominated engagement, together with a marked shortening of transmembrane helices (TM) 5 and 6, has been proposed to explain both ACKR1 broad ligand tolerance and its inability to recruit canonical transducers, with the conformational plasticity of the N-terminal tail allowing on-demand positioning of different chemokines without remodeling of the transmembrane core^13,14^. This plasticity is fine-tuned by sulfation of two N-terminal tyrosines, Tyr30 and Tyr41, whose contribution is both site-and ligand-specific: mutating Tyr41 abolishes binding of CCL2, CCL5, and CXCL1 but not CXCL8, while mutating Tyr30 abolishes CXCL8 binding^15^. This site-specific logic is consistent with the broader principle, established across several chemokine receptors, that N-terminal sulfotyrosines tune chemokine affinity and selectivity in a position-dependent manner^16–18^. Recent structures refine this picture: sulfation at Tyr30/Tyr41 is functionally required for both CCL7 and CXCL8 binding, yet sulfated Tyr41 is clearly resolved and forms a defined ionic-contact network only in the CXCL8-bound structure, remaining structurally elusive in the CCL7 complex despite being equally necessary there^14^. ACKR1’s promiscuity therefore appears to arise not from a single universal binding solution, but from a flexible N-terminal scaffold whose sulfation state is differentially read by distinct chemokines.

On the pathogen side, *P. vivax* Duffy-binding protein (PvDBP) and *S. aureus* leukocidin LukE converge on an overlapping but mechanistically distinct strategy, clamping onto a short, sulfotyrosine-containing segment of the same N-terminal domain: PvDBP forms a receptor-induced dimer that engages an amphipathic helix encompassing residues 19-30 together with sulfated Tyr41, while LukE recognizes the same sulfated Tyr41 through a dedicated sulfotyrosine-binding pocket conserved across several leukocidin S-components^9,15,19,20^. Functional mutagenesis reinforces this convergence: substituting Tyr41, but not the neighboring Tyr30, abrogates both PvDBP-and LukE-mediated cell targeting^11,15,21^. ACKR1 therefore appears to read out a compact sulfotyrosine code in its N-terminus, with host chemokines and pathogen ligands each selecting a different combination of the same one or two sulfated residues to engage the receptor.

The leukocidin heterodimer HlgAB, however, appears to diverge from the logic that governs other ACKR1 ligands. Unlike LukED, HlgAB-induced hemolysis is only weakly competed by CXCL8 and is comparatively insensitive to competition by PvDBP or by antibodies targeting the ACKR1 N-terminus^11^. Whether this reflects a distinct mode of Tyr41 recognition, an additional binding determinant beyond the amino terminus, or some combination of the two remains unresolved. Adding to this uncertainty, we previously showed that the S-component HlgA competes directly with endogenous chemokines for the ACKR1 extracellular domain and that toxin binding allosterically remodels the receptor’s intracellular G protein binding region. Unexpectedly, we also found that the F-component HlgB binds ACKR1 as well^22^, raising the question of whether both subunits, not HlgA alone, contribute to the receptor’s recognition of HlgAB. This question was sharpened by subsequent work showing that HlgB can bind the erythrocyte surface independently of ACKR1 and of HlgA^23^, leaving open how HlgB’s interaction with the receptor relates to this ACKR1-independent binding. Given that the *hlgABC* locus is part of the *S. aureus* core genome, present in over 99% of clinically sequenced strains^24^, HlgAB represents one of the most consistently expressed ACKR1-targeting leukocidins, and one whose structural basis of receptor recognition remains the least resolved piece of ACKR1’s ligand-promiscuity puzzle.

Here, we combine native mass spectrometry, hydrogen-deuterium exchange mass spectrometry, molecular dynamics simulations, small-angle X-ray scattering and ensemble analysis, cell-based assays, and cryo-EM to dissect the molecular basis for HlgA and HlgB recognition by ACKR1. We show that HlgA and HlgB bind the sulfated N-terminus of ACKR1 with distinct affinities and site hierarchies, with a preferred, structurally specific site present in HlgA but absent in HlgB, while both toxins engage additional lower-affinity dynamic sites. We further demonstrate that although the sulfated N-terminus enhances local toxin concentration at the receptor surface, it is dispensable for pore formation, and that productive HlgAB engagement instead requires the extracellular vestibule of ACKR1. Together, these findings reveal a hierarchical, multi-site recognition mechanism that underlies the functional divergence between the S-and F-components of leukocidins in targeting ACKR1.

## Results

### HlgA and HlgB bind to ACKR1 N-terminus with different affinities

To explore the interaction between the N-terminal part of ACKR1 to leukotoxins HlgA and HlgB, native mass spectrometry (nMS) experiments were performed in order to identify direct binding and corresponding stoichiometry, if present. HlgA showed specific binding when mixed with equimolar amounts of ACKR1 sulfated N-terminal peptide ^36^DSFPDY(SO_3_^-^)GANLE^46^ spanning Tyr41 (hereafter ^sulfo^P) with up to two copies of peptide per toxin (Figure 1A). No binding was observed when HlgA was incubated with the non-sulfated version of the peptide (hereafter P) (Supplementary Figure 1A). Interestingly, binding was also observed with a scrambled form of the same sulfated peptide (hereafter *scr*^sulfo^P), in which the sulfated tyrosine was kept highly accessible at the beginning of the sequence. This aligns well with our previous observation that the main driving force behind ACKR1 N-terminal recognition is the sulfotyrosine side chain itself, rather than its specific position within the sequence, with the negative charge of the sulfate group contributing to binding at the positively charged surface of the leukotoxin LukE^20^. For HlgB, specific binding to only one copy was observed when the toxin was incubated at equimolar amount with ^sulfo^P, but to a lesser degree than HlgA, suggesting a lower overall affinity (Figure 1B).

**Figure 1.**
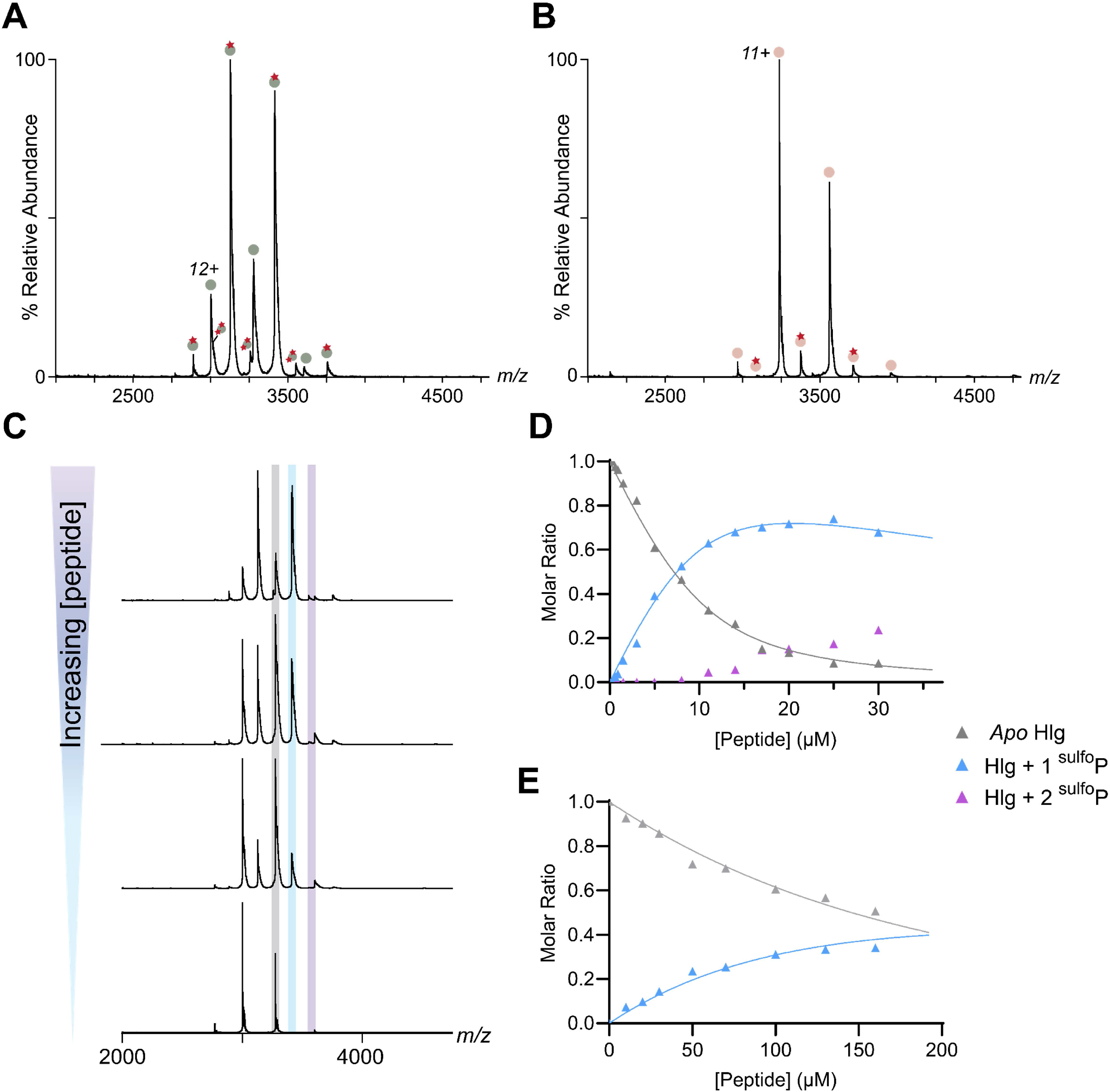
Native MS of sulfated N-terminal ACKR1 binding to HlgA and HlgB. Mixtures of 10 μM ^sulfo^P with 10 μM **(A)** HlgA or **(B)** HlgB, showing binding of two ^sulfo^P to HlgA and binding of one ^sulfo^P to HlgB. Sage circles represent *apo* HlgA at 36 017 ± 1 Da; binding of one and two ^sulfo^P is represented by single and double red stars, respectively, for distributions corresponding to 37 538 ± 1 and 39 053 ± 7 Da. Pink circles represent *apo* HlgB at 35 604 ± 1 Da; binding of one ^sulfo^P is represented by a single red stars corresponding to 37 124 ± 1 Da. **(C)** Representative mass spectra from the titration of 10 μM HlgA with ^sulfo^P, showing the *apo* spectrum (bottom) and spectra obtained upon addition of increasing peptide concentrations (2, 3.5 and 10 μM respectively, bottom to top). The 11^+^ charge state of *apo* HlgA is highlighted in grey, HlgA bound to one peptide in blue, and HlgA bound to two peptides in purple. Representative plots of the mole fraction for **(D)** HlgA *apo* or bound to one or two peptides, and for **(E)** HlgB *apo* or bound to one peptide, determined from a single titration series (dots), with fits from a sequential ligand-binding model (solid lines).

In order to further quantify the differences in the binding of both toxins to ACKR1 N-terminus, we performed a titration using nMS by recording spectra at equilibrium with increasing amounts of peptides added to either HlgA or HlgB (Figure 1C-E). The use of nMS enables the quantification of individual binding events, unlike classical biophysical approaches that measure average binding affinities between a protein and its ligand. In the case of HlgA, the apparent dissociation constant for the binding of the first ^sulfo^P (K_D1_) was of 1.6 ± 0.74 μM (n=4) (Figure 1D) and the binding of the *scr*^sulfo^P had an apparent K_D1_ of 3.22 ± 0.92 μM (n=4) (Supplementary Figure 1B). This demonstrates a similar affinity of both peptides to HlgA, confirming that the main driving force behind ACKR1 N-terminus binding to HlgA is the sulfated Tyr41. We could not quantify binding of the second copy of ^sulfo^P to HlgA, as the combination of overlapping signals with the *apo* peaks and limited resolution at high peptide concentrations kept us from reaching the saturating concentrations needed. Of note, at the highest peptide concentrations tested, an additional third binding signal was observed but considered of very low affinity and/or non-specific under our tested conditions. For HlgB (Figure 1E), the apparent affinity for the ^sulfo^P was nearly 200-fold lower, with K_D1_ = 213 ± 20 μM (n=4). Here also, additional binding of up to two peptides were visible at very high ligand concentration, more likely to be of very low affinity and/or non-specific. Overall, the nMS data indicate that interaction with the N-terminus of ACKR1 is predominantly driven by the sulfated Tyr41. Specifically, HlgA exhibits one higher-affinity and one lower affinity binding event, whereas HlgB predominantly displays a single weak binding event.

### Distinct binding sites of sulfated N-terminal ACKR1 on HlgA and HlgB

In order to identify peptide binding sites on both toxins, we performed differential Hydrogen-Deuterium exchange coupled to MS (ΔHDX-MS) experiments in the presence of the different peptides. Because nMS measurements indicated a micromolar-range affinity, we adjusted the concentrations of the added peptides to ensure sufficient protein occupancy^25^. Accordingly, for a final toxin concentration of 1 µM in the labelling mixture, ^sulfo^P was added at increasing concentrations (2, 20, 75 µM), while *scr*^sulfo^P was added only at the highest concentration (75 µM) (Figure 2). The total number of detected peptides resulting from toxin digestion in all analyzed states was lower compared to that observed in the *apo* state, likely due to ion suppression effects arising from the high concentration of added peptide ligands. Nevertheless, ΔHDX-MS analysis of combined states yielded 51 and 68 peptides, with an overall sequence coverage of 83.7 % and 88.8 % and an average redundancy of 2.78 and 3.08 for HlgA and HlgB respectively (Supplementary Data files 1 and 2). To exclude nonspecific effects of the high peptide concentrations used, we included the non-sulfated control P, which does not bind either toxin, as confirmed by nMS (Supplementary Figure 1A). No significant protection was observed at the sites of interest with P (Figure 2A, B), confirming that protection seen with the sulfated peptides reflects specific toxin-ligand interactions.

**Figure 2.**
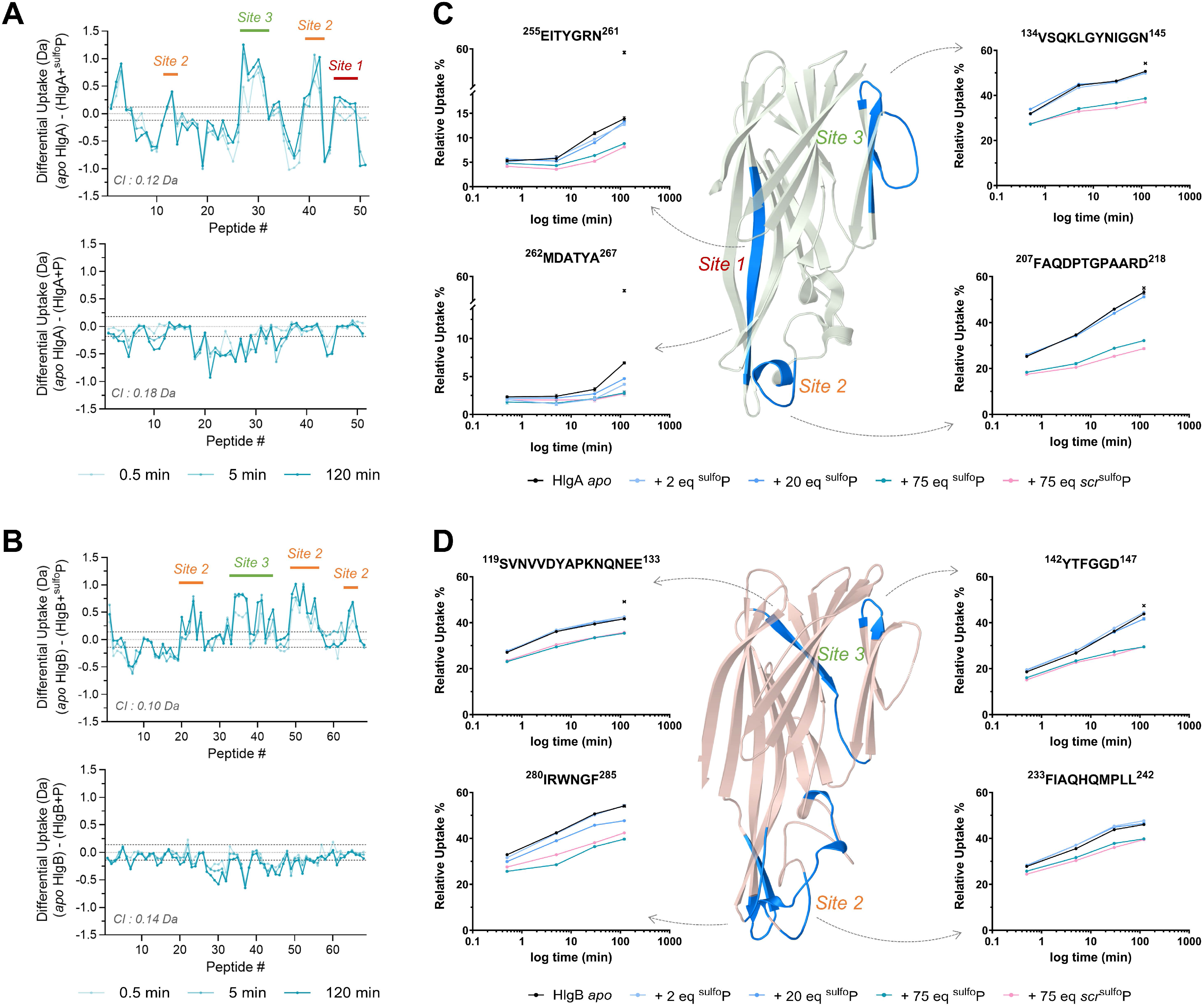
Binding sites of sulfated N-terminal ACKR1 on HlgA and HlgB identified by HDX-MS. Butterfly plots showing differences in average deuterium uptake (ΔHDX) in Da for **(A)** HlgA and **(B)** HlgB across three time points (0.5, 5 and 120 min), comparing the ligand-bound and *apo* states. Top: ΔHDX in the presence of 75 molar equivalent ^sulfo^P; bottom: ΔHDX in the presence of 75 molar equivalent P. Identified peptides of each toxin are arranged along the x-axis from N-to C-terminus. Positive values indicate reduced deuteration (protection); negative values indicate increased deuteration (destabilization) upon ligand addition. Data represent means of triplicate measurements. Dotted grey lines mark the 99% confidence interval threshold for statistically significant ΔHDX. For corresponding coverage and heatmaps, see Supplementary Figures 3 and 4. Deuterium uptake plots for selected peptides of **(C)** HlgA and **(D)** HlgB, mapped onto the corresponding structures (PDB 2QK7 and 2LKF for HlgA and HlgB, respectively) to highlight the main regions protected in the presence of ^sulfo^P. Uptake is shown under the following conditions: ligand-free (black), 2 molar equivalents ^sulfo^P (light blue), 20 molar equivalents ^sulfo^P (medium blue), 75 molar equivalents ^sulfo^P (dark blue), and 75 molar equivalents *scr*^sulfo^P (pink). Maximum deuteration is indicated in asterisk. The sequence of the corresponding residues is shown above each selected peptide. Uptake plots represent the mean ± SD of three technical replicates.

A reduced deuterium uptake in the presence of ^sulfo^P or *scr*^sulfo^P was visible at three main sites on HlgA (Figure 2A, C and Supplementary Figures 2 and 3): site 1, corresponding to the middle of the β-pleated blade 2, between the cap domain and the rim domain; site 2, at the level of the divergent regions in the rim domain; and site 3, at level of the pre-stem domain. At the lowest ligand concentration tested with ^sulfo^P, significant protection was observed at site 1, whereas sites 2 and 3 did not exhibit statistically significant changes in deuterium uptake. Importantly, site 1 is located within a structured β-sheet region that is intrinsically less prone to deuterium exchange compared to the more flexible, solvent-exposed unstructured regions comprising sites 2 and 3. The observation of marked protection at site 1 despite its structured nature suggests a specific and higher-affinity ligand interaction at this site rather than a nonspecific stabilization effect. In contrast, sites 2 and 3 exhibited significant protection only at the highest ligand concentration tested, consistent with comparatively lower-affinity interactions at these more flexible regions. Regarding HlgB, protection was mainly visible at the level of the divergent regions in the rim domain (site 2), spanning multiple peptides, and some protection was observed at the pre-stem level (site 3) (Figure 2B, D and Supplementary Figure 4). This protection was mainly visible at the higher peptide concentrations added, correlating with a relatively higher K_D_ observed for ^sulfo^P binding to HlgB, as compared to HlgA. In contrast to HlgA, no protection was observed at the level of site 1 on HlgB. Altogether, these HDX-MS data outline a hierarchy of ACKR1 peptide binding sites on HlgA, with site 1 behaving as a potential higher-affinity, structurally specific interaction and sites 2 and 3 as lower-affinity, more dynamic contacts. The absence of any protection at site 1 on HlgB indicates that this site is not conserved between the two toxins, consistent with the comparatively lower affinity of HlgB for the ACKR1 peptide.

**Figure 3.**
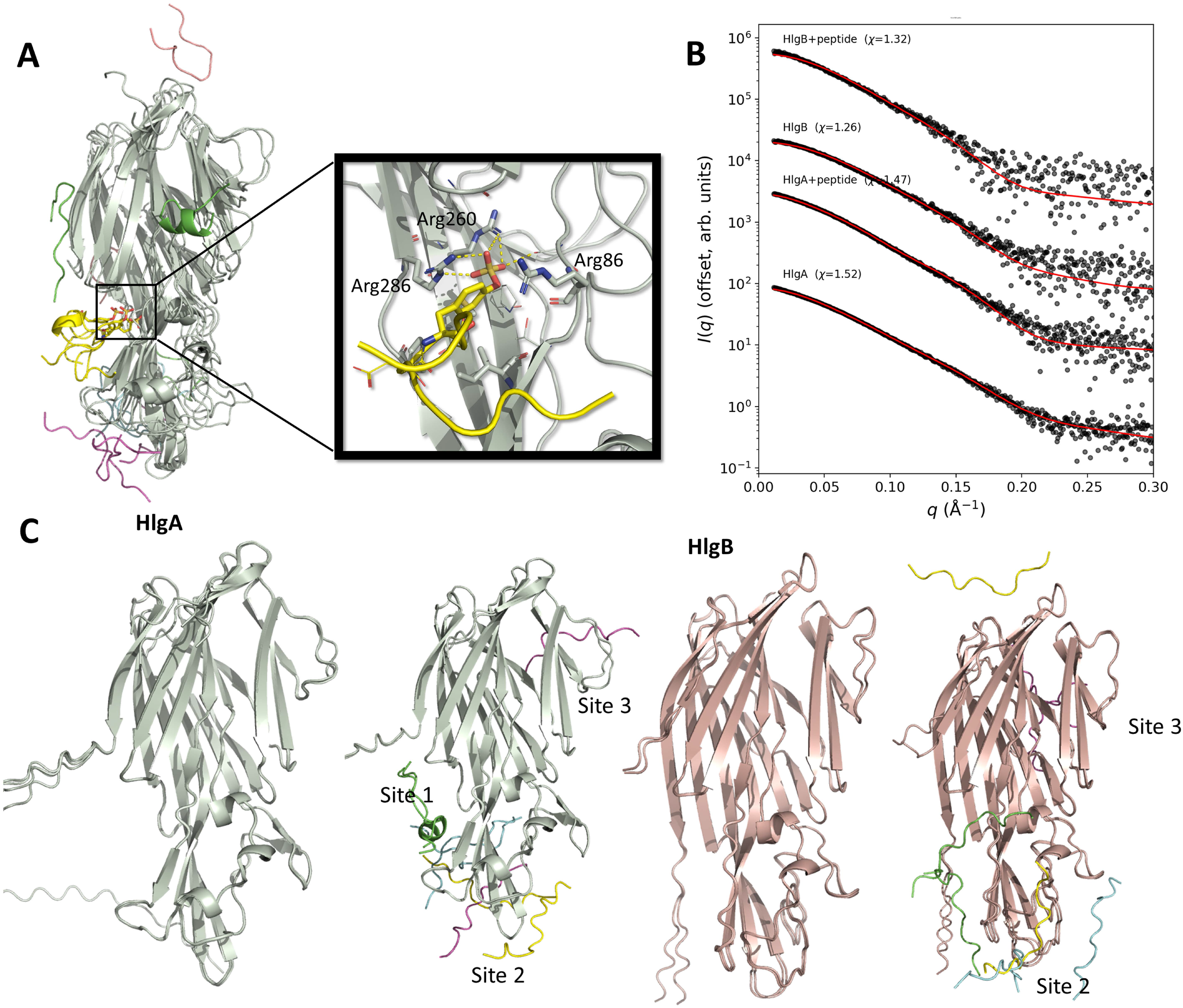
Molecular dynamics and SAXS analysis of sulfated N-terminal ACKR1 binding on HlgA and HlgB. **(A)** Superimposed snapshots of HlgA in presence of 5 copies of ^sulfo^P extracted from MD trajectory at t = 0, 1 and 2 µs. Inset: zoom on the sulfotyrosine binding site 1 at t = 2 µs, highlighting the interactions between the sulphate group and arginine residues 86, 260 and 286 from HlgA. **(B)** Ensemble optimization method (EOM)-fitted SAXS profiles of HlgA and HlgB in the absence of presence of 100 molar equivalents of ^sulfo^P. Experimental data is shown as black spheres and fits are shown as red lines. **(C)** Selected ensembles reproducing the experimental scattering pattern, corresponding to the fits shown in **(B)**, for HlgA, HlgA + ^sulfo^P, HlgB, and HlgB + ^sulfo^P. For HlgA, the C-terminal purification tag is partly omitted for clarity.

### Hierarchical binding of HlgA and HlgB to the N-terminus of ACKR1

To further probe the potential binding locations and mechanism of ACKR1 peptide engagement on HlgA and HlgB, we performed molecular dynamics simulations of each toxin in the presence of five copies of ^sulfo^P, positioned using Chai-1 predictions^26^. For each toxin, a single ∼2 µs trajectory was generated. For HlgA, most peptides showed large conformational changes and/or dynamic binding and unbinding over the course of the simulation (Figure 3A and Supplementary Figure 5A), with one notable exception: the peptide occupying the previously described sulfotyrosine binding site 1 remained associated throughout, with the sulfotyrosine acting as a stable anchor even as the rest of the peptide retained some mobility (Figure 3A). For HlgB, by contrast, all five peptides exhibited dynamic binding and unbinding without evidence of a stably anchored site, consistent with the overall lower affinity of this toxin for ^sulfo^P (Supplementary Figure 5B, C). In both toxins, transient binding events were also observed at sites 2 and 3, in agreement with the HDX-MS data.

To complement these simulations, we used small-angle X-ray scattering (SAXS) to characterize each toxin alone and in the presence of a 100-fold molar excess of ^sulfo^P, with excess free peptide matched by buffer subtraction to isolate the scattering signal of the pure complexes. The data showed no signs of aggregation, aside from a minor low-Q curvature in the HlgA curves that did not affect Guinier analysis or ensemble optimization method (EOM) ensemble fitting^27^ (Supplementary Figure 5D). Kratky plots were relatively similar for both toxins with and without added peptides, suggesting no major changes in overall flexibility (Supplementary Figure 5E). Guinier analysis of HlgA yielded a radius of gyration (Rg) of 2.90 nm and a volume of correlation (Vc)-based molecular weight (MW) estimate of 36.6 kDa (theoretical MW of 36 kDa) (Supplementary Table 1). In the presence of ^sulfo^P, HlgA showed a comparable Rg of 2.84 nm with a MW estimate of 43 kDa, against a theoretical MW of 42 kDa for the toxin bound to four peptides. HlgB alone had an Rg of 2.63 nm, with a MW estimate of 30.3 kDa (theoretical MW of 35.6 kDa); in the presence of ^sulfo^P, its Rg increased to 3.02 nm and its MW estimate to 37.6 kDa, against a theoretical MW of 41.6 kDa for the toxin bound to three peptides (Supplementary Table 1, Figure 3B and Supplementary Figure 5D, E). The difference in Rg between the two apo toxins reflects their different purification tags (His-tag for HlgB, His-and Twin-Strep-tag for HlgA, included to improve purification and solubility) rather than a difference in overall fold. The estimated MW are within the errors of SAXS measurements and consistent with monomeric HlgA and HlgB binding several copies of ^sulfo^P.

To determine the peptide-binding stoichiometry underlying these scattering profiles, we generated structural ensembles (50 seeds × 5 models = 250 models per condition) using Protenix v2^28^, modeling the apo form of each toxin alongside complexes with one, two, three or four bound peptides, and fitted each ensemble against the corresponding experimental SAXS data using the EOM fitting. Comparison of the resulting χ² values (Supplementary Table 1) showed that the best fits for both toxins were obtained with three to four peptides bound, rather than with lower-occupancy models. For HlgA, the selected ensembles placed two peptides at site 1, with additional peptides occupying sites 2 and 3 (Figure 3C); for HlgB, bound peptides were positioned mainly at site 2 and occasionally site 3, with no occupancy at site 1 (Figure 3C). These stoichiometries and site assignments are consistent with the MS data described above. Collectively, the MD simulations and SAXS-ensemble analysis reveal that HlgA and HlgB exploit fundamentally different binding site hierarchies: site 1 is occupied and structurally stable only in HlgA, whereas both toxins engage the more dynamic sites 2 and 3, highlighting the functional divergence between S-and F-components in their recognition of ACKR1.

### Site 1 in HlgA is the primary binding site for ACKR1 sulfated Tyr41

Taken together, the ΔHDX results combined with apparent K_D_ measurements by nMS, molecular dynamics simulations, and SAXS-based ensemble analysis, suggest that site 1 is the high-affinity binding site for ^sulfo^P at the level of HlgA. Structural comparison of site 1 between HlgA and HlgB reveals important differences underlying their distinct binding affinities (Figure 4A, B). While two of the three arginines forming site 1 are conserved between both toxins (Arg260/Arg273 and Arg286/Arg298), the third arginine is not: Arg86 in HlgA is replaced by a glycine at the equivalent position in HlgB (Gly88), reducing the positive charge density at the binding site. More strikingly, HlgB contains an additional β-strand not present in HlgA that forms a salt bridge between HlgB Glu322 and the equivalent of Arg286 (Arg298), effectively neutralizing this critical positive charge and explaining why site 1 is functionally ablated in HlgB. To validate the functional importance of site 1, we generated a mutant of HlgA in which we disrupted the two arginines most critical for differential binding between the two toxins: Arg86 and Arg286 were mutated to glycine and glutamic acid, respectively (hereafter HlgA_Δsite1_). We spared Arg260 from mutation because it remains conserved in HlgB despite the overall ablation of site 1 in that toxin, and because the structural analysis suggests that the Arg86→Gly substitution and the Glu322-Arg298 salt bridge in HlgB are the primary mechanisms responsible for site 1 inactivation. nMS titration of ^sulfo^P showed a decreased binding affinity to HlgA_Δsite1_ (K_D1_ = 81 ± 12 μM, n=3) (Figure 4C), and ΔHDX revealed protection at the level of sites 2 and 3, whereas the protection at site 1 observed with wild-type HlgA was lost in the HlgA_Δsite1_ mutant (Figure 4D, E and Supplementary Figure 6). Altogether, these results demonstrate that although both HlgA and HlgB bind to the N-terminal sulfated region of ACKR1, HlgA engages site 1 with higher affinity, whereas interactions at sites 2 and 3 on both proteins exhibit an overall low affinity.

**Figure 4.**
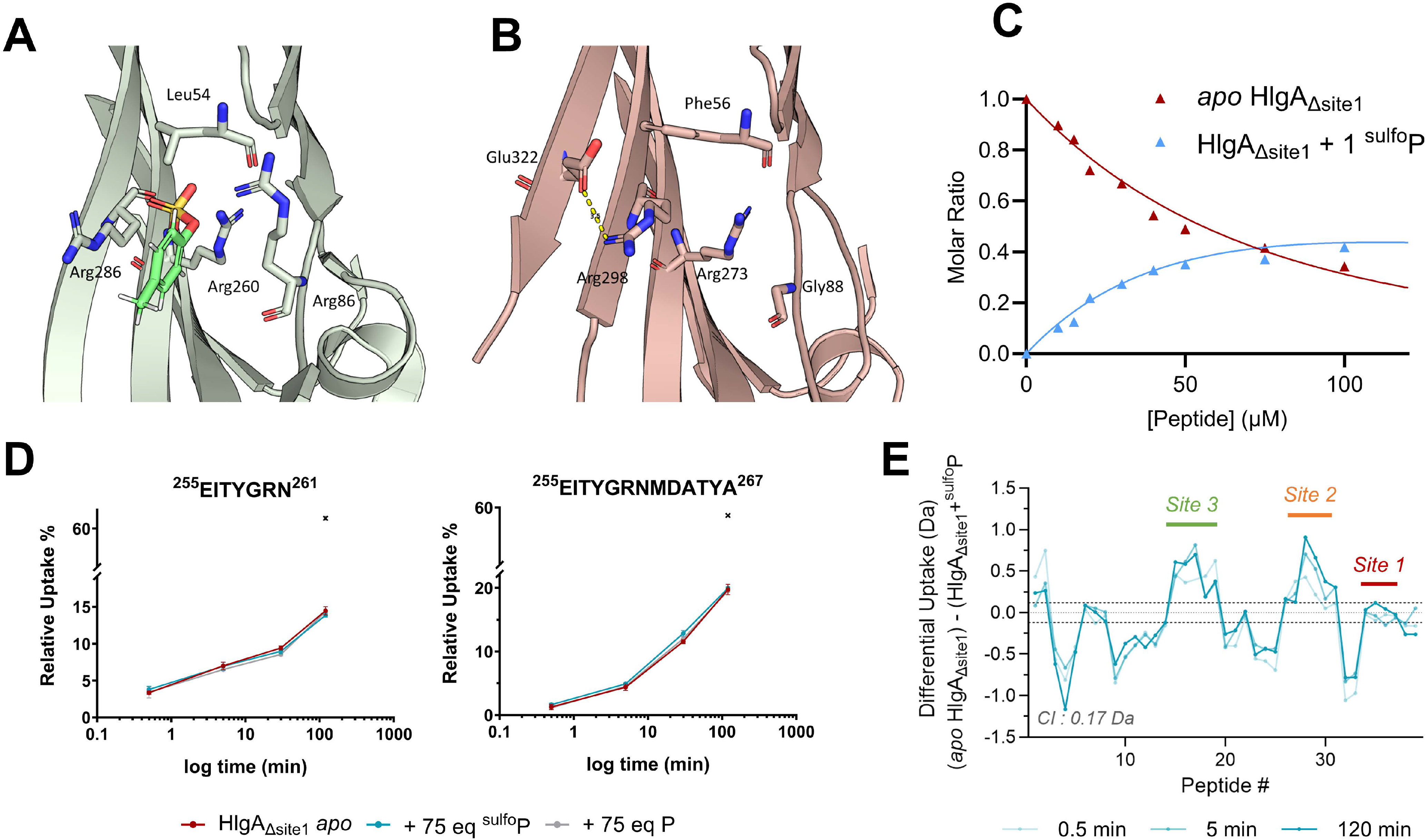
Primary binding site for sulfated Tyr41 on HlgA. Structural comparison of site 1 between **(A)** HlgA and **(B)** HlgB. Modeled sulfotyrosine binding to HlgA is presented in green. **(C)** nMS titration of 10 μM HlgA_Δsite1_ with ^sulfo^P, showing the extracted molar ratio of the *apo* protein and the protein bound to one peptide as a function of increasing ^sulfo^P concentration. Dots represent experimental data points from a single titration series; solid lines represent fits to a sequential ligand-binding model used to determine the apparent K_D_. **(D)** Deuterium uptake plots for peptides identified at site 1, under the following conditions: ligand-free (burgundy), 75 equivalents ^sulfo^P (blue), and 75 equivalents P (grey). Maximum deuteration is indicated in asterisk. The sequence of the corresponding residues is shown above each selected peptide. Uptake plots represent the mean ± SD of three technical replicates. **(E)** Butterfly plots showing differences in average deuterium uptake (ΔHDX) in Da for HlgA_Δsite1_ across three time points (0.5, 5 and 120 min), comparing the ligand-bound (75 molar equivalent of ^sulfo^P) and *apo* states. Identified peptides of each toxin are arranged along the x-axis from N-to C-terminus. Positive values indicate reduced deuteration (protection); negative values indicate increased deuteration (destabilization) upon ligand addition. Data represent means of triplicate measurements. Dotted grey lines mark the 99% confidence interval threshold for statistically significant ΔHDX. For corresponding coverage and heatmaps, see Supplementary Figure 6.

### Binding to the N-terminus of ACKR1 is not mandatory for pore formation

In order to explore the role of toxins binding to the sulfated N-terminus of ACKR1 in pore formation, we monitored pore formation in HEK293 cells by tracking cytosolic Ca^2+^ concentration (fluorescence) that increases rapidly and substantially upon adding HlgAB^29^, but only in cells over-expressing a cognate receptor (Supplementary Figure 7 A-D). Indeed, a spike in Ca^2+^ influx was visible upon adding HlgAB in a concentration-dependent manner, only in cells over-expressing wild-type ACKR1 but not in mock cells or in cells over-expressing a control class A G protein-coupled receptor (μ-opioid receptor, µOR). To further validate that the observed increased intracellular Ca^2+^ concentration is dependent on pore-formation by HlgAB and not on the activation of other receptor-dependent pathways, we used dominant-negative variants of both toxins (hereafter δHlg). These variants are impaired in pore formation as they lack the glycine-rich region in the pre-stem^30^. While competitive inhibition of d2-CCL5 binding to Tb-ACKR1 by time-resolved fluorescence resonance energy transfer (TR-FRET) clearly shows that both δHlgA and δHlgB maintain their binding capacity to ACKR1 (Supplementary Figure 7E), adding a mixture of δHlgAB did not induce any increased intracellular Ca^2+^ levels in the presence of ACKR1 (Supplementary Figure 7F). This confirms that pore formation is indeed impaired with the δHlgAB variants and that increased Ca^2+^ levels observed with the wild-type HlgAB in the presence of ACKR1 is related to pore formation.

To compare between the different conditions and biological replicates, we monitored the maximum-to-minimum fluorescence obtained upon increasing HlgAB concentration in the presence of full-length ACKR1 for wild-type HlgA compared to HlgA_Δsite1_ (Figure 5A and Supplementary Figure 7 A, G). Mutation of site 1 slightly increased the EC50 for Ca^2+^ influx, thereby decreasing the potency of pore formation. To further explore this, we performed competitive inhibition of d2-CCL5 binding to Tb-ACKR1 by TR-FRET using both wild-type HlgA and HlgA_Δsite1_ (Figure 5B). The results show that, while the IC50 was similar for both proteins, the maximal d2-CCL5 displacement achieved by HlgA_Δsite1_ was decreased by nearly 50% compared to the wild-type HlgA (Figure 5B). This demonstrates that mutating site 1 on HlgA impairs its ability to fully displace CCL5 but does not fully prevent the toxin to bind the receptor. The binding data thus indicate the existence of at least two different binding sites of HlgA on ACKR1. Finally, when deleting residues ^36^DSFPDY(SO_3_^-^)GANLE^46^ from the N-terminus of ACKR1 (hereafter ACKR1_ΔNter_), both HlgAB and HlgA_Δsite1_B were still capable of forming pores with similar potencies, indicating that the contribution of HlgA site 1 to pore-formation depends on the sulfated N-terminus. However, EC50 in the presence of ACKR1_ΔNter_ was increased compared to wild-type receptor (Figure 5A and Supplementary Figure 7 H, I). Altogether, these data demonstrate that HlgA occupies at least two binding sites on ACKR1, and that the N-terminal part of ACKR1 encompassing sulfated Tyr41, while not mandatory for pore formation, enhances pore-formation potency, likely by increasing the local concentration of toxin at the receptor surface.

**Figure 5.**
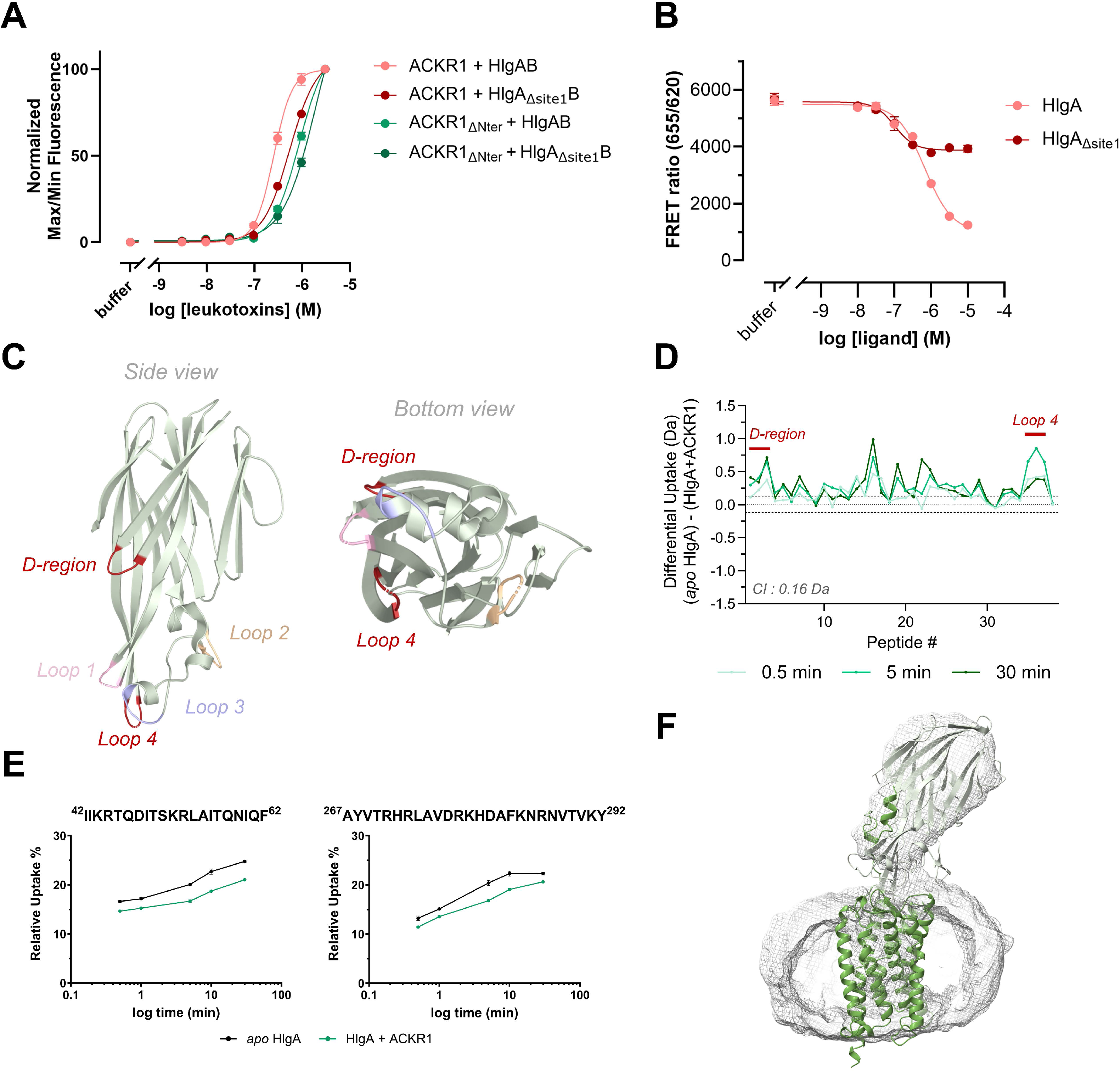
Pore formation requires the engagement with the extracellular vestibule of ACKR1. **(A)** Normalized maximum-to-minimum (Max/Min) Ca^2+^ influx data in HEK293 cells overexpressing wild-type ACKR1 or ACKR1_ΔNter_ in the presence of increasing concentrations of HlgAB or HlgA_Δsite1_B. Data shown are the mean ± SD of one experiment performed in duplicates and are representative of three independent experiments. EC50 values, representing the average ± SD of three independent experiments, are as follows: HlgAB in the presence of ACKR1, 0.30 ± 0.08 μM; HlgA_Δsite1_B in the presence of ACKR1, 0.57 ± 0.12 μM; HlgAB in the presence of ACKR1_ΔNter_,1.21 ± 0.30 μM; and HlgA_Δsite1_B in the presence of ACKR1_ΔNter,_ 1.32 ± 0.55 μM. **(B)** Competitive TR-FRET showing the displacement of d2-CCL5 from Tb-ACKR1 in the presence of increasing amounts of wild-type HlgA or HlgA _Δsite1_. Data shown are the mean ± SD of one experiment performed in duplicates and are representative of three independent experiments. IC50 values, representing the average ± SD of three independent experiments, are as follows: HlgA, 064 ± 0.17 μM; and HlgA_Δsite1_, 0.11 ± 0.02 μM. **(C)** Cartoon representation of HlgA (PDB 2QK7) showing the loop structures in the rim domain and the D-region in the cap domain. **(D)** Butterfly plots showing differences in average deuterium uptake (ΔHDX) in Da for HlgA across three time points (0.5, 5 and 30 min), comparing the ACKR1-bound and *apo* states. Identified peptides of each toxin are arranged along the x-axis from N-to C-terminus. Positive values indicate reduced deuteration (protection); negative values indicate increased deuteration (destabilization) upon ACKR1 addition. Data represent means of triplicate measurements. Dotted grey lines mark the 99% confidence interval threshold for statistically significant ΔHDX. For corresponding coverage and heatmap, see Supplementary Figure 8. **(E)** Deuterium uptake plots for peptides identified at the D-region and Loop 4 under in the apo state (black) and the ACKR1-bound state (green). The sequence of the corresponding residues is shown above each selected peptide. Uptake plots represent the mean ± SD of three technical replicates. **(F)** Flexible fitting of the Protenix model into the low-resolution experimental cryo-EM map using iSOLDE, shown as cartoon superimposed on the cryo-EM density (gray mesh), demonstrating agreement between the predicted atomic model and the experimental electron density.

### The extracellular vestibule of ACKR1 is required for toxin productive engagement

Since the N-terminus of ACKR1 is not required for pore formation in living cells but instead enhances pore-formation potency by increasing local toxin concentration, we next examined the effect of full-length ACKR1 on the global deuteration profile of HlgA *in vitro*, to identify additional toxin regions influenced by the full-length receptor but not by its N-terminal peptide alone. Full-length ACKR1 was produced in *Sf9* cells, purified in n-dodecyl-d-maltoside (DDM) detergent, and incubated with HlgA at equimolar amounts. ΔHDX-MS analysis comparing *apo* and receptor-bound HlgA revealed protection at most ^sulfo^P-protected sites, albeit to a lesser degree (ΔRFU ∼5%, Supplementary Figure 8). This reduced magnitude reflects a technical limitation: unlike with the peptide, the receptor-to-toxin ratio could not be increased to compensate for the low affinity of the interaction. Full-length ACKR1 protected two regions previously shown to be implicated in HlgAB hemolytic activity that were not protected by ^sulfo^P (Figure 5C-E and Supplementary Figure 8). First, ACKR1 protected loop 4 in the divergent region of the HlgA rim, previously shown to be critical for hemolysis, in contrast to loops 1 and 2, protected by ^sulfo^P, which instead assist HlgA binding to erythrocytes^31^. This suggests ^sulfo^P-binding regions serve mainly to concentrate toxin at the membrane, not to drive hemolysis *per se*. Second, ACKR1, unlike ^sulfo^P, protected the D-region established as pivotal for HlgAB hemolytic activity^31,32^, underscoring the importance of the full-length receptor for HlgAB-mediated hemolysis.

To further resolve the HlgA-ACKR1 binding mode, we performed cryo-EM analysis on a mixture of the two proteins. 2D classification showed a density corresponding to HlgA atop a micelle containing the receptor (Supplementary Table 2 and Supplementary Figure 9), with class averages revealing multiple toxin orientations relative to the receptor, consistent with conformational dynamics within the complex. Interestingly, the toxin consistently engaged one of the receptor’s top edges. Because ACKR1 produced in *Sf9* cells is not uniformly sulfated at Tyr41^14^, this heterogeneity likely contributed to the conformational variability and reduced stability of the complex. This dynamic behavior limited the resolution to ∼5.8 Å; nonetheless, the cryo-EM density clearly defines the complex’s overall architecture (Figure 5F), revealing the global organization and relative positioning of the receptor and toxin rather than supporting residue-level interpretation. HlgA consistently bound the receptor at an angle of ∼65° across reconstructions (Supplementary Figure 10). Structural models generated with ProteniX v2^28^ showed a good overall fit to the experimental cryo-EM density and were refined in ISOLDE^33^ to further optimize map agreement (Figure 5E and Supplementary Figure 11). The generated model shows extensive engagement between ACKR1 N-terminus and HlgA, along with contacts between the toxin and the receptor’s extracellular region (Supplementary Figure 11 and Supplementary Table 3). HlgA regions that showed protection from deuterium exchange in the presence of ^sulfo^P and the full-length receptor made multiple contacts with the ACKR1 N-terminus, consistent with HDX-MS data. In addition, loop 4 in the divergent region of HlgA formed several contacts with the extracellular vestibule of ACKR1, engaging the extracellular loop 2 (ECL2) and the orthosteric pocket. Together, the cryo-EM reconstruction, fitted models, and HDX-MS data converge on a shared interaction architecture, supporting a two-step model in which the sulfotyrosine site capture the toxins at the receptor surface, while pore formation additionally requires vestibule engagement mainly via HlgA loop 4.

## Discussion

Together, our data support a model in which ACKR1 presents two spatially and functionally uncoupled binding surfaces to the HlgAB bicomponent leukotoxin (Figure 6). Its sulfated N-terminal Tyr41 is recognized by the toxin heterodimer through a hierarchy of affinities. Loops in the divergent region of the rim domain and the pre-stem region of both subunits offer only low affinity contacts to sulfo-Tyr41, whereas a single predominant site for this same residue exists in HlgA alone, positioned in the middle of β pleated blade 2 between the cap and rim domains (site 1). Mechanistically, site 1 higher kinetic stability likely reflects coordination of sulfo-Tyr41 by a specific binding pocket, including 3 arginine guanidinium groups and several hydrophobic residues interacting with the Tyr41 aromatic ring, substantially slowing off rate relative to the lower affinity sites. The corresponding site is not conserved in HlgB, which the receptor N-terminus therefore engages only weakly. Yet ACKR1’s sulfated N-terminus is not required to trigger pore formation by HlgAB. Deleting or mutating site 1 does not abolish pore formation, and HlgAB retains comparable potency in blocking CCL5 binding to ACKR1 regardless of which toxin subunit carries the site 1^22^. This dissociation between N-terminal affinity and pore forming capacity indicates that ACKR1’s sulfo-Tyr41 acts as a capturing site, concentrating toxins locally at the cell surface, rather than as a trigger for the conformational transitions underlying oligomerization and membrane insertion. Productive pore formation instead requires a second, functionally decisive engagement, with the receptor’s extracellular vestibule, as demonstrated by our cell-based, HDX-MS and cryo-EM data. Mainly, the ECL2 and the orthosteric pocket of ACKR1 contact loop 4 of HlgA, previously shown to be required for hemolysis by HlgAB^31^. This two-step recognition architecture, in which ACKR1 N-terminus captures the toxin and its extracellular vestibule triggers pore formation, reconciles several observations that would otherwise appear contradictory, and clarifies why HlgA and HlgB, despite sharing a receptor, behave as functionally asymmetric partners.

**Figure 6.**
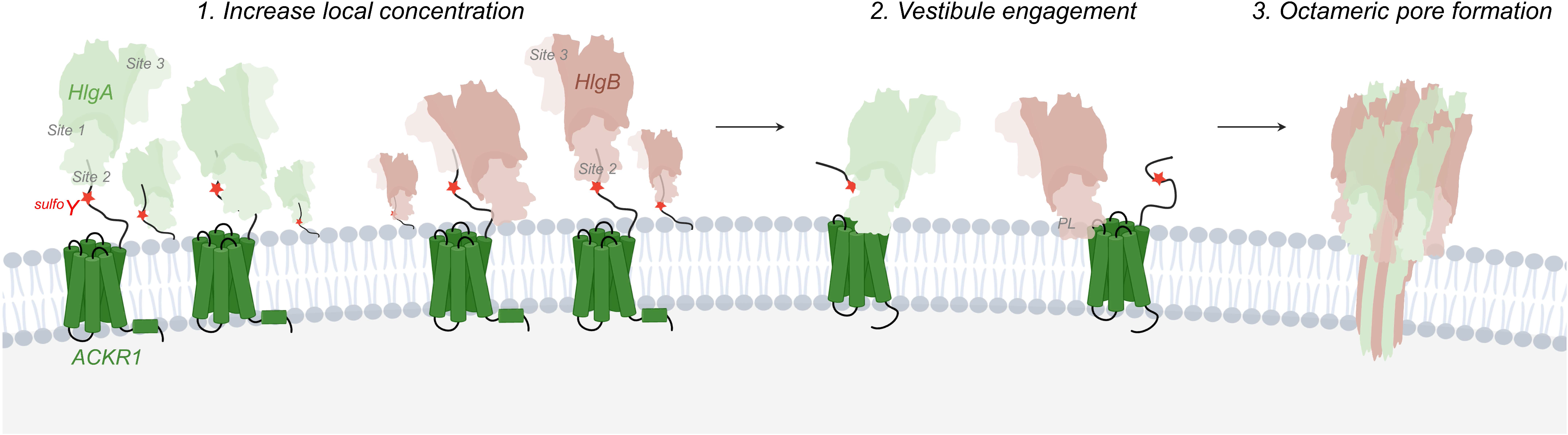
First steps of toxin interactions with ACKR1 leading to pore formation. Interaction of HlgA and HlgB with the sulfated N-terminus of ACKR1 increases local toxin concentration (step 1). This step is not sufficient to trigger pore formation. Engagement with the vestibule of the receptor (step 2) and HlgB binding to membrane phospholipids (PL) leads to octameric pore formation.

ACKR1 itself appears to be engaged differently by the different toxin pairs that converge on it. Our finding that HlgAB requires the receptor extracellular vestibule for productive engagement, rather than the amino terminus alone, aligns with and extends previous mutagenesis data showing that ACKR1 determinants differentially affect HlgAB and LukED pore-formation^11^. Mutation of the disulfide forming Cys51 selectively impaired LukED but not HlgAB, while mutations in ECL2 (Glu202Ala) and ECL3 (Asp283Ala) had opposing effects on the two toxin pairs. Together with our structural data placing HlgA’s loop 4 against the ECL2 and the orthosteric pocket of ACKR1, this indicates that ACKR1 offers largely non overlapping surfaces to HlgAB and LukED despite being a shared target. This is consistent with the loop swap incompatibility reported by Peng *et al.*^31^, in which HlgA loop 4 could not functionally substitute for LukE loop 4 and vice versa. In this light, vestibule engagement is best understood as a receptor-guided pre-step to oligomerization, orienting the captured toxins toward the membrane.

This framework helps reconcile a recurring theme across previously published and our current work, namely that ACKR1 apparent contribution shrinks as toxin concentration rises, for reasons that likely differ depending on which of these two steps is being bypassed. Spaan *et al.*^11^ found Tyr41 important for hemolysis by HlgAB, seemingly at odds with our finding that the high affinity site is dispensable for pore formation. We suspect this reflects a concentration effect specific to the capture step. Consistent with this, Peng *et al.*^31^ showed that mutating Tyr41 required substantially more toxin to reach full hemolysis, suggesting that ACKR1 N-terminus accelerates lysis under limiting toxin concentrations by locally concentrating toxin, without being strictly required once sufficient toxin reaches the vestibule by mass action alone. A similar logic may explain the recent report that HlgB binds erythrocytes independently of HlgA or ACKR1^23^. The HlgB rim domain harbors an intrinsic phosphocholine binding site (Trp202, Glu217, Arg223) that directly engages choline headgroups in the plasma membrane domain^34–36^, offering a plausible, receptor independent basis for this membrane association. At the elevated toxin concentrations used in such assays, this might reflect a membrane association driven by excess toxin, rather than engagement through the normal, receptor-templated capture and orientation pathway. The *in vitro* structural studies of the formed octameric HlgAB pore raise a related but distinct point: whether pore-formation was triggered in liposomes^37^ or by 2-methyl-2,4-pentanediol^38^, the toxin concentrations used in both cases were far above the physiological range. Because these systems lack a receptor entirely, pore formation there cannot rely on either capture or vestibule mediated orientation, and instead likely reflects mass action compensating for both steps simultaneously, a scenario with no clear physiological counterpart given that pore formation is essentially receptor-dependent^39^.

Viewed alongside its other ligands, ACKR1 achieves promiscuity not through one flexible binding mode, but by allocating different combinations of a limited set of surfaces, the N-terminal sulfotyrosine code, the extracellular vestibule, and the broader ECL/TM region, to different ligand classes. This distributed use of the receptor surface, rather than a single adaptable interface, is a structural basis for ACKR1 promiscuity, and predicts that different ligand classes will remain differentially sensitive to mutations at each site.

## Methods

### Peptides and protein constructs

Peptides were custom synthesized at the SynBio3 facility (University of Montpellier), using solid phase peptide synthesis, at >99% purity. Protein constructs were custom made by Genecust. All peptide sequences and protein constructs used in this study are listed in Supplementary Table 4.

### HlgA and HlgB production and purification

Toxins were expressed in BL21 (DE3) *E. coli* cells (New England Biolabs). Transformed cells were grown at 37°C in Luria–Bertani broth for both toxins supplemented with 100 mg/mL ampicillin or kanamycin to an optical density of 0.6. Expression was then induced at 20 °C by addition of 0.5 mM isopropyl ß-D-1-thiogalactopyranoside. Cells were harvested by centrifugation (3,000 rpm), and cell pellets were purified. Cells were first lysed by sonication in a lysis buffer consisting of 20 mM Tris (pH 8), 300 mM NaCl, 2 mg/mL iodoacetamide (Sigma-Aldrich), and protease inhibitors (Leupeptin [Euromedex], Benzamidine, and PMSF [Sigma-Aldrich]). Lysed cells were centrifuged (16,000 rpm), and the supernatant was purified using affinity chromatography. For HlgA, the supernatant was loaded on a Strep-Tactin® Superflow® 50% suspension (IBA Lifesciences). The resin was washed with 4 column volumes (CV) of buffer 1 consisting of 50 mM Tris pH 7.5 and 1 M NaCl, and 2 CV of buffer 2 consisting of 50 mM Tris pH 7.5 and 300 mM NaCl. Bound HlgA was eluted with buffer 2 supplemented with 2.5 mM Desthiobiotin (IBA Lifesciences). For HlgB, δHlgA, and δHlgB, the supernatant was loaded on Ni-NTA resin (ThermoFisher). The resin was washed with 10 CV of buffer 1 and with 10 CV buffer 2 supplemented with 40 mM imidazole. Bound toxins were eluted with buffer 2 supplemented with 400 mM imidazole. The eluted solution of all toxins was concentrated to 500 μL using 10-kDa spin filters (Millipore) and further purified by SEC on a Superdex 75 Increase 10/300 column (GE Healthcare) in buffer 2. Fractions containing monodisperse toxins were collected, concentrated, flash-frozen and kept at -80°C.

### Native MS acquisition and analysis

Prior to nMS analysis, toxins were dialyzed for 24 to 48 h in 200 mM ammonium acetate buffer pH 8 (Sigma-Aldrich) to desalt and buffer exchange the samples. Prior to nMS analysis, mixtures of 9 or 10 µM toxins were preincubated with the different peptides for 30 min to reach equilibrium, at different molar ratio (1:0.1 to 1:50). nMS spectra were recorded on a Synapt G2-Si high-definition mass spectrometer (HDMS) (Waters Corporation) using MassLynx 4.2 software (Waters Corporation). The measurements were conducted in positive resolution mode. Samples were introduced into the ion source using borosilicate emitters (Thermo Fisher Scientific). Optimized instrument parameters were as follows: capillary voltage 1.4 kV, sampling cone voltage 120 V, offset voltage 70 V, transfer collision voltage 5 V, trap collision energy 5V, and argon flow rate 7 mL/min. Native MS spectra were deconvoluted using UniDec^40^ and the intensities of toxin and toxin-ligand species were converted to mole fraction for each peptide concentration added, with Y_0_ corresponding to *apo*, Y_1_ to singly-bound and Y_2_ to doubly-bound toxins. Apparent equilibrium dissociation constants were determined as described in^41^; for our data, a sequential binding model of up to two ligands gave the best fits, and only the first dissociation constant is reported, as the titration range did not reach saturation for the second binding event, leaving K_D2_ poorly constrained. For every analyzed condition, three to four independent titration series were performed and the reported K_D_ values represent the average ± SD of all titration series performed.

### HDX-MS data acquisition and analysis

HDX-MS experiments were performed using a Synapt G2-Si HDMS coupled to nanoAQUITY ultra–high-performance liquid chromatography (UPLC) with HDX Automation technology (Waters Corporation). Toxins were concentrated up to 20 µM, and optimization of the sequence coverage was performed on undeuterated controls. Mixtures of toxins with the different peptides (2, 20 and 75 molar equivalents) were preincubated 15 min prior to HDX-MS analysis. Analysis of freshly prepared toxin *apo* and toxin peptide mixtures were performed as follows: 3 µL sample are diluted in 57 µL undeuterated buffer for the reference or deuterated buffer 2 for other timepoints. The final percentage of deuterium in the deuterated buffer was 95%. Deuteration was performed at 15°C for 30, 300, 1800, and 7200 s with the peptides, and for 30, 60, 300, 600 and 1800 s with ACKR1. Maximum deuteration was performed from denaturated toxin to correct back exchange as previously described ^42^. 15 µL of toxin dried in SpeedVac to replace SEC buffer with 15µL of 7M guanidine, then heated at 90°C for 5 min. 285 µL of labeling buffer was added to it, and then heated for 10 min at 50°C. Next, 50 µL of the reaction sample is quenched in 50 µL quench buffer (KH_2_PO_4_ 50 mM and K_2_HPO_4_ 50 mM, pH 2.3) at 0 ° C for 30 sec. A total of 80 µL quenched sample is loaded onto a 50-µL loop and injected on an Enzymate pepsin column (300 Å, 5 mm, 2.1 mm × 30 mm, Waters Corporation) maintained at 15°C, with 0.2% formic acid at a flowrate of 100 mL/min and an additional backing pressure of 6,000 psi controlled by the HDX regulator kit (Waters Corporation). The peptides are then trapped at 0°C on a Vanguard column (ACQUITY UPLC BEH C18 VanGuard Precolumn, 130 Å, 1.7 mm, 2.1 mm × 5 mm, Waters Corporation) for 3 min before being loaded at 40 µL/min onto an Acquity UPLC column (ACQUITY UPLC BEH C18 Column, 1.7 mm, 1 mm × 100 mm, Waters Corporation) kept at 0 ° C. Peptides are subsequently eluted with a linear gradient (0.2% formic acid in acetonitrile solvent at 5% up to 35% during the first 6 min, then up to 40% and 95% over 1 min each) and ionized directly by electrospray on a Synapt G2-Si mass spectrometer (Waters Corporation). HDMSE data were obtained by 20-to 30-V trap collision energy ramp. Lock mass accuracy correction was made using a mixture of leucine enkephalin and Glu-fibrinopeptide B. The pepsin column was then washed three times with Guanidine-HCl 1.5 M, acetonitrile 4%, and formic acid 0.4%, and a blank is performed between each sample in order to minimize the carryover. All time points were performed in triplicates. Peptide identification was performed from undeuterated data using the ProteinLynx global Server (version 3.0.3, Waters Corporation). Peptides are filtered by DynamX (version 3.0, Waters Corporation) using the following parameters: minimum intensity of 1,000, minimum product per amino acid of 0.2, maximum error for threshold of 10 ppm. All peptides were manually checked, and data were curated using DynamX. Statistical analysis of all ΔHDX data was performed using Deuteros 2.0^43^, and only peptides with a 99% confidence interval were considered. HDX summary tables and data tables can be found in Supplementary Data files 1 and 2. Of note, several regions of HlgA showed increased deuterium uptake in the presence of ^sulfo^P, *scr*^sulfo^P, and the control P. Because this destabilization occurred with the non-binding control as well, it is unlikely to reflect specific binding effects. Possible explanations include nonspecific, buffer-related factors, such as counterion or salt differences introduced with the synthetic peptides and/or minor synthesis impurities.

### Molecular dynamics simulations

Molecular dynamics (MD) simulations were performed using GROMACS^44^ with the CHARMM36 force field. System setup was performed using CHARMM-GUI^45^, which generated topologies for the HlgA–^sulfo^P and HlgB–^sulfo^P complexes. Five copies of ^sulfo^P were positioned around each toxin based on Chai-1 predictions^26^. The systems were solvated in a cubic box of TIP3P water with 0.15 M NaCl, energy-minimized, and equilibrated using the standard CHARMM-GUI protocol (NVT followed by NPT). Production runs were carried out at 300 K and 1 bar for approximately 2 µs per system with a 2 fs time step, and coordinates were saved every 1 ns for analysis. RMSD of toxins and peptide chains were calculated using GROMACS routines after alignment of CA atoms onto the toxin.

### Small-angle X-ray scattering (SAXS) data collection and analysis

SAXS measurements were performed at the BM29 beamline at the European Synchrotron Radiation Facility (ESRF) at room temperature. HlgA and HlgB (20 µM) were measured alone and in the presence of a 100-fold molar excess of ^sulfo^P. Excess free peptide was matched by buffer subtraction to isolate the scattering signal of the protein– peptide complexes. Data were processed and merged using PRIMUS^46^. Guinier analysis was performed in the low-Q region to determine the radius of gyration (Rg), and molecular weight (MW) was estimated using the volume of correlation (Vc) method^47^. Kratky plots were generated to assess changes in overall flexibility between apo and peptide-bound states.

### Ensemble-based structure prediction and SAXS fitting

To determine peptide-binding stoichiometry and architecture, structural ensembles were generated using Protenix v2^28^. For each condition (*apo* toxin and toxins bound to one up to four ^sulfo^P), 250 independent models were generated (50 seeds × 5 models per seed), with peptide poses informed by Chai-1 predictions and the MD simulations. Each ensemble was fitted against the experimental SAXS data using GAJOE^27^, which optimizes the mixture of conformers to reproduce the experimental scattering curve using a genetic algorithm. The goodness of fit was assessed using the χ² metric; the ensemble with the lowest χ² value was selected as the best description of the solution-state ensemble. The final selected ensembles were analyzed to determine peptide occupancy at the three binding sites (sites 1, 2, and 3).

### Ca^2+^ influx experiments in living cells

30 ng of the different receptors were added to 5.10^4^ cells/well for transfection in 96-well back plates. 24 h post-transfection, the cells were incubated for 1 h at 37 °C with 50 µL/well Cal-520 AM (AAT Bioquest) working solution at 1.25 µM in FLEX buffer (Hank’s balanced salt solution, 20 mM HEPES, 1 mM MgSO_4_, 3.3 mM Na_2_CO_3_, 1.3 mM CaCl_2_). After incubation, cells were washed with 130 µL/well with FLEX buffer, and toxins where then injected into each well, at increasing concentrations, approximately 1 min after the start of the readings. Cal-520 fluorescence signals were quantified on a FDSS-µCELL plate reader (Hamamatsu Photonics K.K) at 1.3 sec per reading frame; samples were illuminated at 480 nm and fluorescence was acquired at 540 nm. For data analysis, maximum fluorescence signal was the sum of the signal obtained over 40 readings after toxins injection (130 to 182 sec), whereas the minimum is the sum of the signal obtained over 40 readings before toxins injection (1 to 52 sec). Max/Min was normalized for every condition tested separately. Each toxin concentration was performed in duplicate, and experiments were repeated at least 3 times. Dose-response curves were generated using GraphPad Prism 6® (GraphPad Software, Inc., San Diego, CA).

### Competitive binding assay in living cells by TR-FRET

30 ng of SNAP-ACKR1 were added to 5.10^4^ cells/well for transfection. 24 h post-transfection, the plate was washed twice with TagLite® (Cisbio, Codolet, France) and the extracellular SNAP-receptors were labelled with SNAP-Lumi4-Tb (100 nM, Cisbio, Codolet, France) for 1 h at 37°C. The plate was washed three times with TagLite®. 12 nM of d2-labelled CCL5 and increasing concentration of toxins (δHlgA, δHlgB or an equimolar mixture of δHlgA and δHlgB) were added to cells. Lumi-4-fluorescence signals were observed on a PHERAstar plate reader (BMG Labtech): samples were illuminated at 337 nm and fluorescence was acquired at 620 nm (donor) and 665 nm (TR-FRET) over time. The ratio of the signals (665/620) was plotted versus the toxin concentration. Dose-response curves were generated using GraphPad Prism 6® (GraphPad Software, Inc., San Diego, CA).

### ACKR1 production and purification

Expression of ACKR1 was carried out in Sf9 insect cells (Life Technologies) using the pFastBac baculovirus system (Thermofischer) as previously described^22^. After thawing the frozen cell pellets, cells were lysed by osmotic shock adding 1/10 (v/v) lysis buffer consisting of 10 mM Tris (pH 7.5), 1 mM EDTA and containing 2 mg/mL of iodoacetamide (Sigma) and protease inhibitors (Leupeptin (Euromedex), Benzamidine and PMSF (Sigma)). Lysed cells were centrifuged (16,000 rpm) and the membrane pellets were suspended in a 1/20 (v/v) solubilization buffer consisting of 50 mM HEPES (pH 7.5), 150 mM NaCl, 0.5% (w/v) n-dodecyl-D-maltoside (DDM, Anatrace), 0.1% (w/v) cholesteryl-hemi-succinate (CHS, Sigma), 2 mg/mL of iodoacetamide and protease inhibitors. Receptors were extracted from the membrane pellets using a glass dounce grinder and the extracted mixture was stirred for 1 h at 4 °C, then centrifuged (16,000 rpm). The supernatant was loaded by gravity flow onto anti-Flag M2 antibody resin. The resin was washed with 10 column volumes (CV) of a DDM wash buffer consisting of 50 mM HEPES (pH 7.5), 150 mM NaCl, 0.1% (w/v) DDM and 0.02% (w/v) CHS. The bound receptor was eluted in the wash buffer 2 supplemented with 0.4 mg/mL Flag-peptide. The eluted solution of receptors was concentrated to 500 μL using 50 kDa spin filters (Millipore) and further purified by size exclusion chromatography on a Superdex 200 Increase 10/300 column (GE Healthcare) in the last wash buffer. Fractions containing monodisperse ACKR1 were collected, concentrated, flash-frozen and kept at -80°C.

### Cryo-EM sample preparation and image acquisition

Before grids preparation, freshly prepared ACKR1 at a concentration of 29 µM were mixed with 1.5-fold equimolar ratio with the HlgA toxin. Following 30 min of incubation, 3 µl of the mixture were applied on glow-discharged (25 mA, 10 s) Quantifoil Gold 1.2/1.3 300-mesh holey carbon grids (Quantifoil), blotted 4 s, and then flash-frozen in liquide thane using a Leica EM GP2. Grids were pre-screened on a Glacios microscope (Thermo Scientific) of the EM platform of the Institute of Structural Biology (IBS), Grenoble, France. Cryo-EM data were collected on a Titan Krios G3i microscope (Thermo Fisher Scientific) operating at 300 kV at the CM01 beamline (ESRF, Grenoble, France). Micrographs were collected on a Gatan K3 detector using a Gatan Quantum LS energy filter. Image acquisition was performed in counting mode with gain normalization enabled with EPU software. In total, 29,791 movies were collected at a magnification of 105k, corresponding to a pixel size of 0.84 Å, using an electron dose of 19.2 electrons/pixel/s and a total exposure time of 1.45 s per movie, yielding a total dose of 39.5 electrons/Ų spread over 40 frames. An energy filter slit width of 20 eV was used to remove inelastically scattered electrons. Two acquisitions per hole were performed with defocus values ranging from −1.0 to −2.2 µm in 0.2 µm increments. Data were acquired in TIFF LZW compressed format.

### Cryo-EM data processing and predicted model refinement

Cryo-EM data for the ACKR1-HlgA complex were processed using cryoSPARC (v4.7.1)^48^. Movie frames were aligned with Patch Motion Correction, and Contrast Transfer Function (CTF) parameters were estimated through Patch CTF correction. Automatic Gaussian blob detection (estimated diameter = 120 to 160, elliptical and circular blob) was used for particle picking, and picked particles were extracted with a box size of 294 Å and downscaled to 2.9 Å/pixel. Extracted particles underwent reference-free 2D classification (100 classes). After iterative 2D classification, the best 2D classes were subjected to multiple rounds of ab-initio model reconstruction in 4 or 5 classes. Particles from the best class were re-extracted at a pixel size of 1.26 Å for refinement. After NU-refinement, particles underwent global CTF refinement, reference-based motion correction, and a final NU-Refinement, resulting in a map with an overall resolution of 5.83 Å (FSC 0.143) (Supplementary Figure 9).

The predicted Protenix v2 model of the ACKR1-HlgA complex was flexibly fitted into the low-resolution cryo-EM density map using iSOLDE^33^ employing secondary-structure restraints and intrachain distance restraints to maintain model geometry while allowing for conformational adjustments to fit the experimental electron density. The fitted model and cryo-EM map were visualized using UCSF ChimeraX and PyMOL.

## Data availability

The HDX-MS and native MS data have been deposited to the ProteomeXchange Consortium via the PRIDE^49^ partner repository with the dataset identifier PXD082142. HDX summary tables and data tables can be found in Supplementary Data Tables 1 and 2. The cryo-EM dataset measured at ESRF CM01 is available here: “Bacia, M., Desfosses, A., & Leyrat, C. (2027). Cryo-EM analysis of the atypical chemokine receptor ACKR1 in complex with Staphylococcus aureus pore forming toxin component HlgA. European Synchrotron Radiation Facility. https://doi.org/10.15151/ESRF-ES-1565620501”. The reconstructed map was deposited in OneDep wwPDB and attributed with the EMD-59457 code.

## Supporting information

Suuplementary Information

## Acknowledgments

This work was supported by the Centre National de la Recherche Scientifique (CNRS), Institut National de la Santé et de la Recherche Médicale (INSERM), and the University of Montpellier. We thank Luc Brunel and Pascal Verdié from SynBio fo peptide synthesis. C.B. was supported from Agence Nationale de la Recherche grant LEUKOCEPTOR (ANR-21-CE44-0007). Native and HDX-MS platforms used at the Montpellier Proteomics Platform (PPM, BioCampus) were co-financed by the European Regional Development Fund (ERDF) and the Occitanie region. PPM is a member of the national Proteomics French Infrastructure (ProFI UAR 2048) supported by the French National Research Agency (ANR-24-INBS-0015, Investments for the future F2030). We used the platforms of the Grenoble Instruct-ERIC centre (ISBG; UAR 3518 CNRS-CEA-UGA-EMBL) within the Grenoble Partnership for Structural Biology (PSB), supported by FRISBI (ANR-10-INBS-0005-02) and GRAL, financed within the University Grenoble Alpes graduate school (Ecoles Universitaires de Recherche) CBH-EUR-GS (ANR-17-EURE-0003). The IBS EM facility is supported by the Rhône-Alpes Region, the Fondation Recherche Medicale (FRM), the fonds FEDER and the GIS-Infrastrutures en Biologie Sante et Agronomie (IBISA). We acknowledge the European Synchrotron Radiation Facility (ESRF) for provision of synchrotron radiation facilities under proposal IDs MX-2620 MX-2482 and on beamlines CM01 and BM29. We thank the staff of the ESRF and EMBL Grenoble for assistance and support in using the jointly operated Structural Biology beamlines.

## Contributions

A.F, E.L, Z.A and C.L.D produced and purified the proteins. E.L performed native MS and analyzed HDX-MS experiments. T.G performed and analyzed HDX-MS experiments. I.B and O.O.O. performed and analyzed cell-based assays. A.F., A.D., M.B and C.L performed and analyzed cryo-EM experiments. C.L and R.B.B performed and analyzed MD and SAXS data. S.G and C.B. conceptualized the project, supervised the study and acquired fundings. C.B. wrote the manuscript with input from A.F, C.L and S.G.

