## Supplementary material for "Distinct atypical chemokine receptor 1 determinants underlie bacterial toxins recognition and pore formation": Suuplementary Information

[1]IGF, Univ Montpellier, CNRS, INSERM, 141 rue de la Cardonille, Montpellier, 34094, France.

[2] Institut de Biologie Structurale, Université Grenoble Alpes, CEA, CNRS (IBS), 38044, Grenoble, France.

[#]Contributed equally

### Supplementary figures

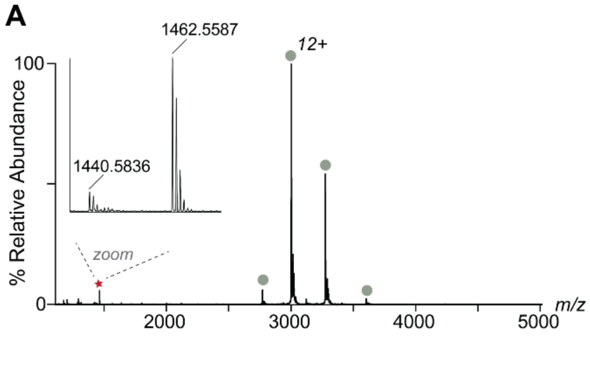

**Supplementary Figure 1. (A)** nMS spectrum of 10 µM HlgA mixed with 10 µM of the non-sulfated N-terminal ACKR1 peptide P, showing no binding to the peptide. Sage circles represent *apo* HlgA at 36 017 ± 1 Da. Inset shows the MH$^+$ of the free peptide at 1440 m/z and the resulting sodium adduct at 1462 m/z (red star). **(B)** Representative plots of the mole fraction for HlgA *apo* or bound to 1 or 2 *scra*$^{sulfo}$P determined from one titration series (dots) and resulting fit from sequential ligand-binding model (solid lines).

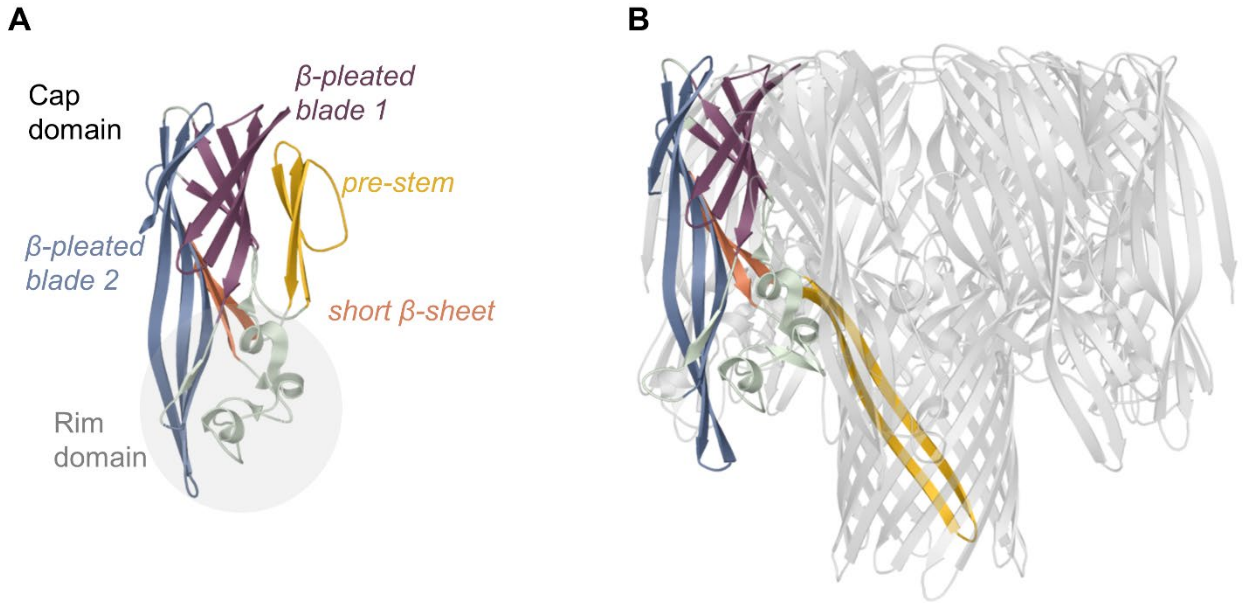

**Supplementary Figure 2.** Cartoon representation of HlgA **(A)** in its monomeric state (PDB 2QK7), and **(B)** in its octameric pore structure formed with HlgB (PBD 3B07). The three different domains of HlgA protomers, similar to the HlgB protomer (not shown), include the β-sandwich cap domain, the pre-stem that forms the transmembrane β-barrel pore in the octameric structure, and a globule rim domain at the bottom (disordered regions and bottom of the of β-pleated blade 2, highlighted by a grey circle). The rim domain contains the divergent loops, and is behind the specificity for receptor recognition of the different leukocidins.

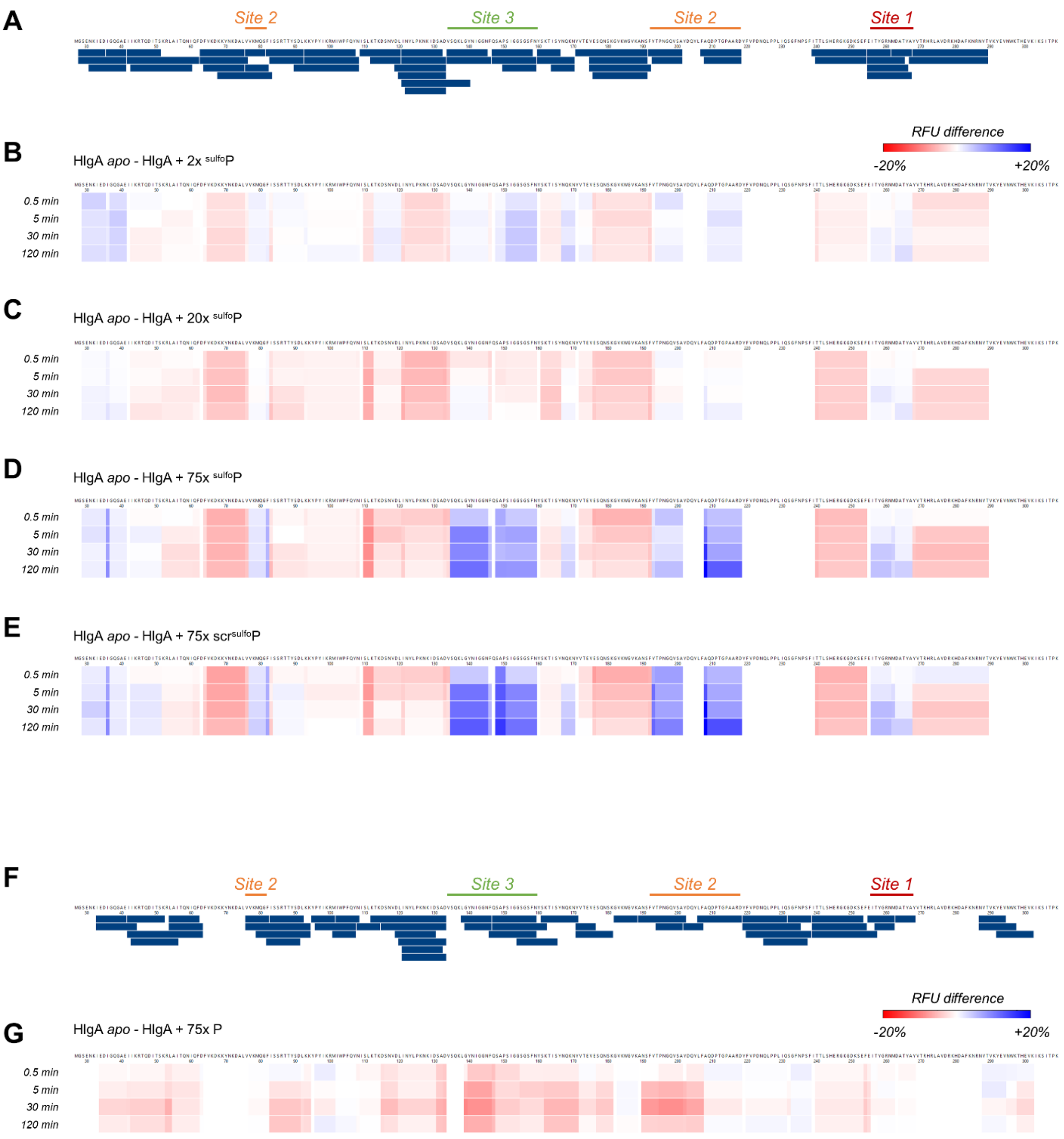

**Supplementary Figure 3. (A)** Coverage map of HlgA showing the common manually curated peptides (blue bars) for all the states analysed in the presence of $^{sulfo}$P and scr $^{sulfo}$P: 51 total peptides detected, representing 83.7% sequence coverage with 2.78 redundancy. The position of the three different binding sites detected on the toxin is labeled above the sequence. **(B)-(E)** Heatmaps showing the percent relative fractional uptake difference (*apo* minus ligand-bound) for HlgA in the presence of 2, 20, or 75 molar equivalents of $^{sulfo}$P, and 75 molar equivalents of *scr*$^{sulfo}$P. **(F)** Coverage map of HlgA showing the common manually curated peptides (blue bars) for *apo* HlgA and HlgA+P: 51 total peptides detected, representing 83.4% sequence coverage with 2.53 redundancy. The position of the three different binding sites detected on the toxin is labeled above the sequence. **(G)** Heatmap showing the percent relative fractional uptake difference (*apo* minus ligand-bound) for HlgA in the presence of 75 molar equivalents of P. Each heatmap shows peptide position (x-axis) versus deuterium exchange time point (y-axis) of 0.5, 5, 30 and 120 min; color indicates the magnitude and direction of uptake difference according to the scale bar (red = increased uptake/deprotection, blue = decreased uptake/protection).

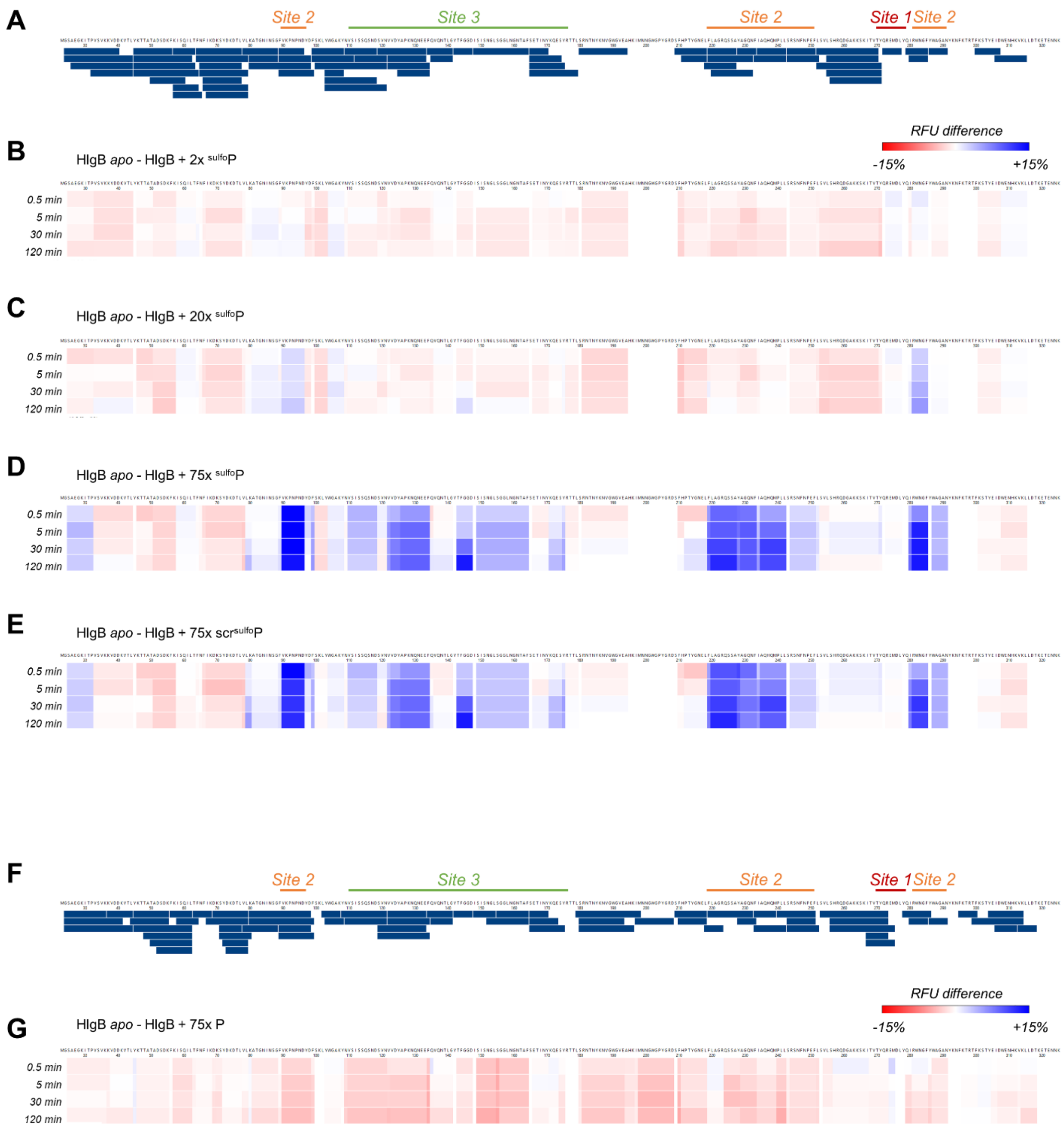

**Supplementary Figure 4. (A)** Coverage map of HlgB showing the common manually curated peptides (blue bars) for all the states analysed in the presence of $^{sulfo}$P and scr $^{sulfo}$P: 68 total peptides detected, representing 88.8% sequence coverage with 3.08 redundancy. The position of the three different binding sites detected on the toxin is labeled above the sequence. **(B)-(E)** Heatmaps showing the percent relative fractional uptake difference (*apo* minus ligand-bound) for HlgB in the presence of 2, 20, or 75 molar equivalents of $^{sulfo}$P, and 75 molar equivalents of *scr*$^{sulfo}$P. **(F)** Coverage map of HlgB showing the common manually curated peptides (blue bars) for *apo* HlgB and HlgB+P: 68 total peptides detected, representing 94.1% sequence coverage with 2.66 redundancy. The position of the three different binding sites detected on the toxin is labeled above the sequence. **(G)** Heatmap showing the percent relative fractional uptake difference (*apo* minus ligand-bound) for HlgB in the presence of 75 molar equivalents of P. Each heatmap shows peptide position (x-axis) versus deuterium exchange time point (y-axis) of 0.5, 5, 30 and 120 min; color indicates the magnitude and direction of uptake difference according to the scale bar (red = increased uptake/deprotection, blue = decreased uptake/protection).

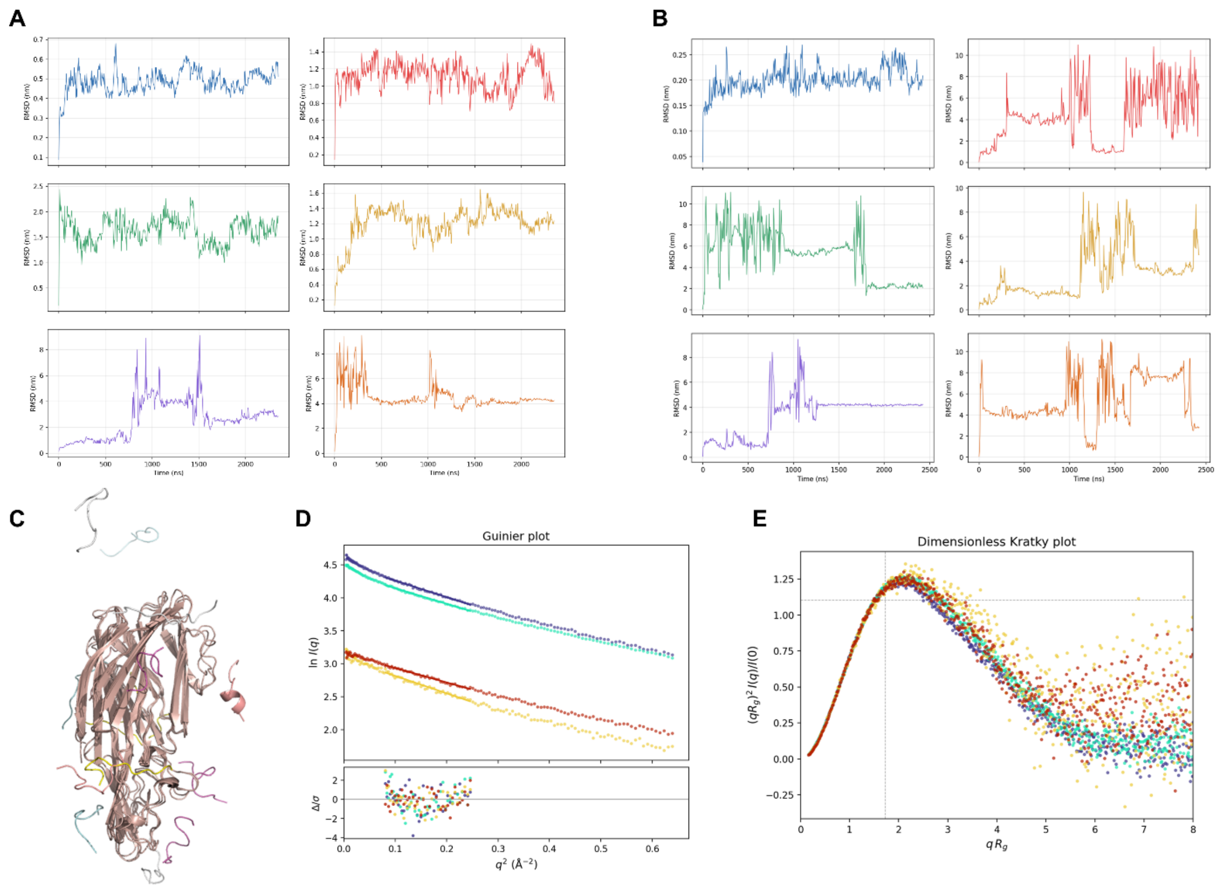

**Supplementary Figure 5**: Cα RMSD analysis of molecular dynamics simulation of **(A)** HlgA and **(B)** HlgB (blue trace), in the presence of five copies of $^{sulfo}$P (red, green, yellow, purple, and orange traces for peptides 1 to 5). **(C)** Superimposed snapshots analysis of molecular dynamics simulation of HlgB in the presence of five copies of $^{sulfo}$P. **(D)** Guinier analysis and **(E)** normalized Kratky plots of measured SAXS data.

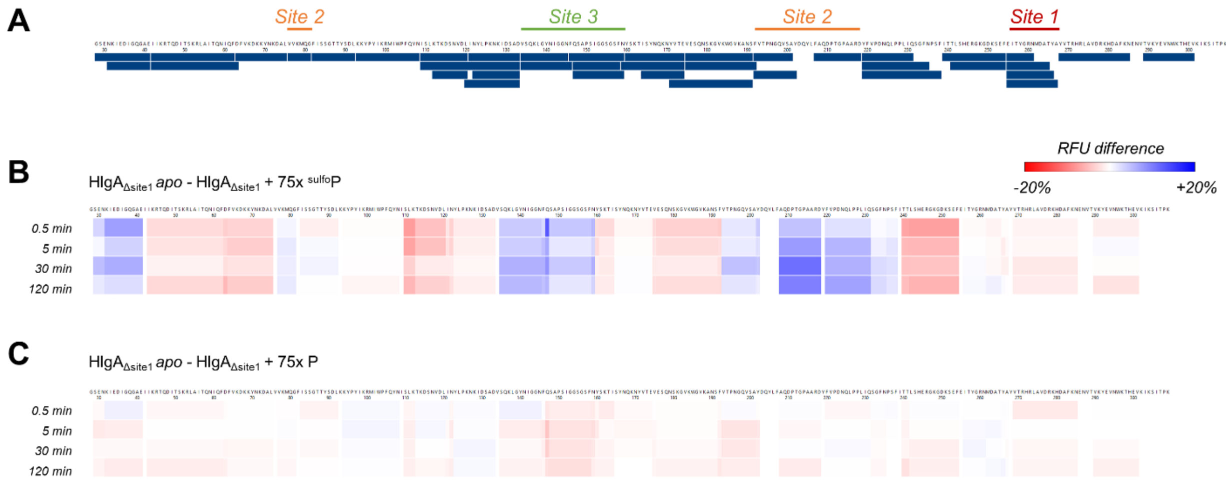

**Supplementary Figure 6. (A)** Coverage map of HlgA$_{\Delta site1}$ showing the common manually curated peptides (blue bars) for all the states analysed in the presence of $^{sulfo}$P and P: 39 total peptides detected, representing 94.7% sequence coverage with 2.05 redundancy. The position of the three different binding sites detected on the wild-type HlgA is labeled above the sequence. **(B)**, **(C)** Heatmaps showing the percent relative fractional uptake difference (*apo* minus ligand-bound) for HlgA in the presence of 75 molar equivalents of $^{sulfo}$P and P. Each heatmap shows peptide position (x-axis) versus deuterium exchange time point (y-axis) of 0.5, 5, 30 and 120 min; color indicates the magnitude and direction of uptake difference according to the scale bar (red = increased uptake/deprotection, blue = decreased uptake/protection).

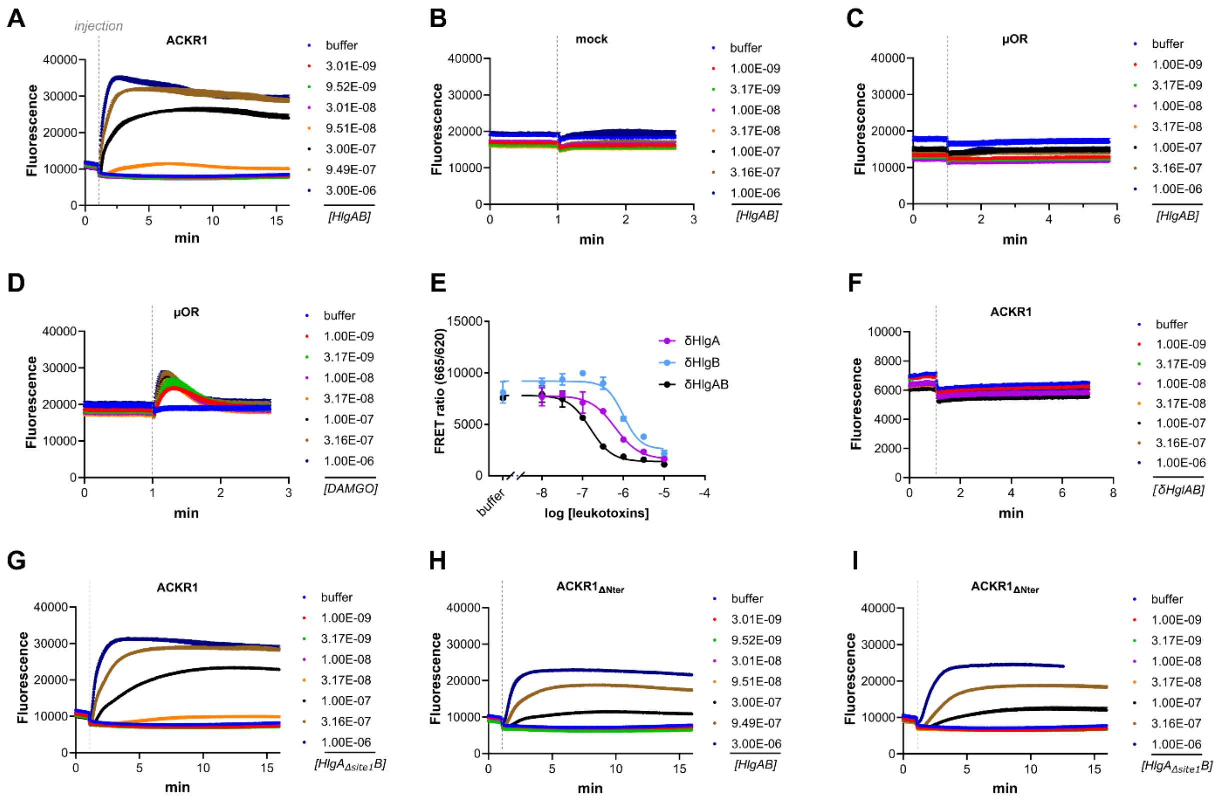

**Supplementary Figure 7. (A-C)** Ca$^{2+}$ influx traces obtained upon addition of increasing concentration of different ligands (HlgAB or DAMGO) in HEK293 cells overexpressing ACKR1, μOR or empty vectors (mock). The injection of the ligands at around 1 min of reading is labeled in dashed line. **(E)** Dose-response curves showing the decrease in TR-FRET ratio between Tb-ACKR1 and d2-CCL5 upon addition of δHlgA, δHlgB and δHlgAB equimolar mixture. Data shown are the mean +/- SEM of one experiment performed in triplicates and are representative of 2 independent experiments. IC50 values: δHlgA 0.60 ± 0.04 μM, δHlgB 1.58 ± 0.83 μM, δHlgAB 0.14 ± 0.02 μM. Values represents the average +/- SD of two independent experiments performed in triplicates. **(F-I)** Ca$^{2+}$ influx traces obtained upon addition of increasing concentration of different ligands (δHlgAB, HlgA$_{\Delta site1}$B and HlgAB) in HEK293 cells overexpressing ACKR1 or ACKR1$_{\Delta Nter}$. The injection of the ligands at ~1 min of reading is labeled in dashed line.

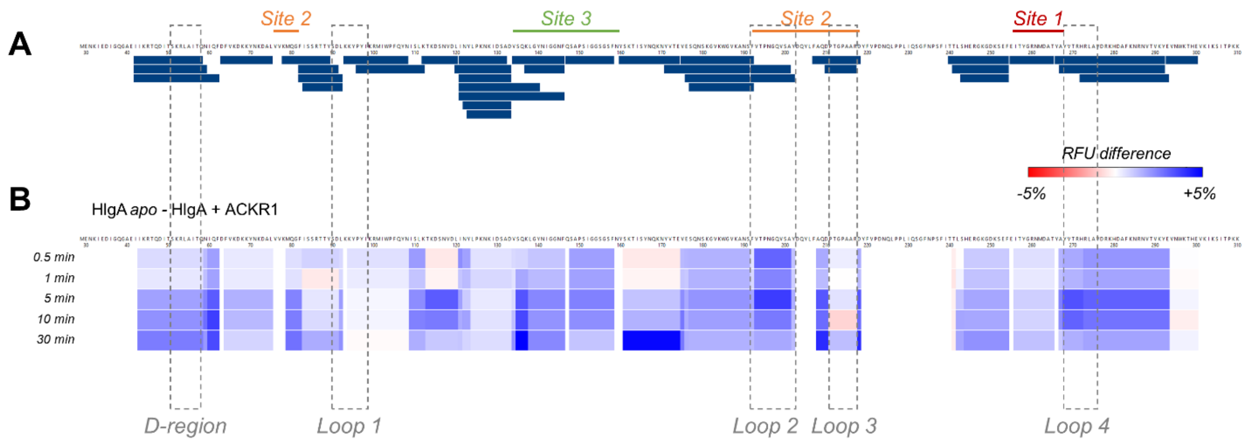

**Supplementary Figure 8. (A)** Coverage map of HlgA showing the common manually curated peptides (blue bars) for all the states analysed in the presence of ACKR1: 38 total peptides detected, representing 81.9% sequence coverage with 2.42 redundancy. The position of the three different binding sites (1, 2 and 3) detected on the wild-type HlgA in the presence of $^{sulfo}$P is labeled above the sequence. The position of the different loop regions and the D-region are indicated in dashed rectangles. **(B)** Heatmap showing the percent relative fractional uptake difference (*apo* minus ACKR1-bound) for HlgA. The heatmap shows peptide position (x-axis) versus deuterium exchange time point (y-axis) of 0.5, 1, 5, 10 and 30 min; color indicates the magnitude and direction of uptake difference according to the scale bar (red = increased uptake/deprotection, blue = decreased uptake/protection).

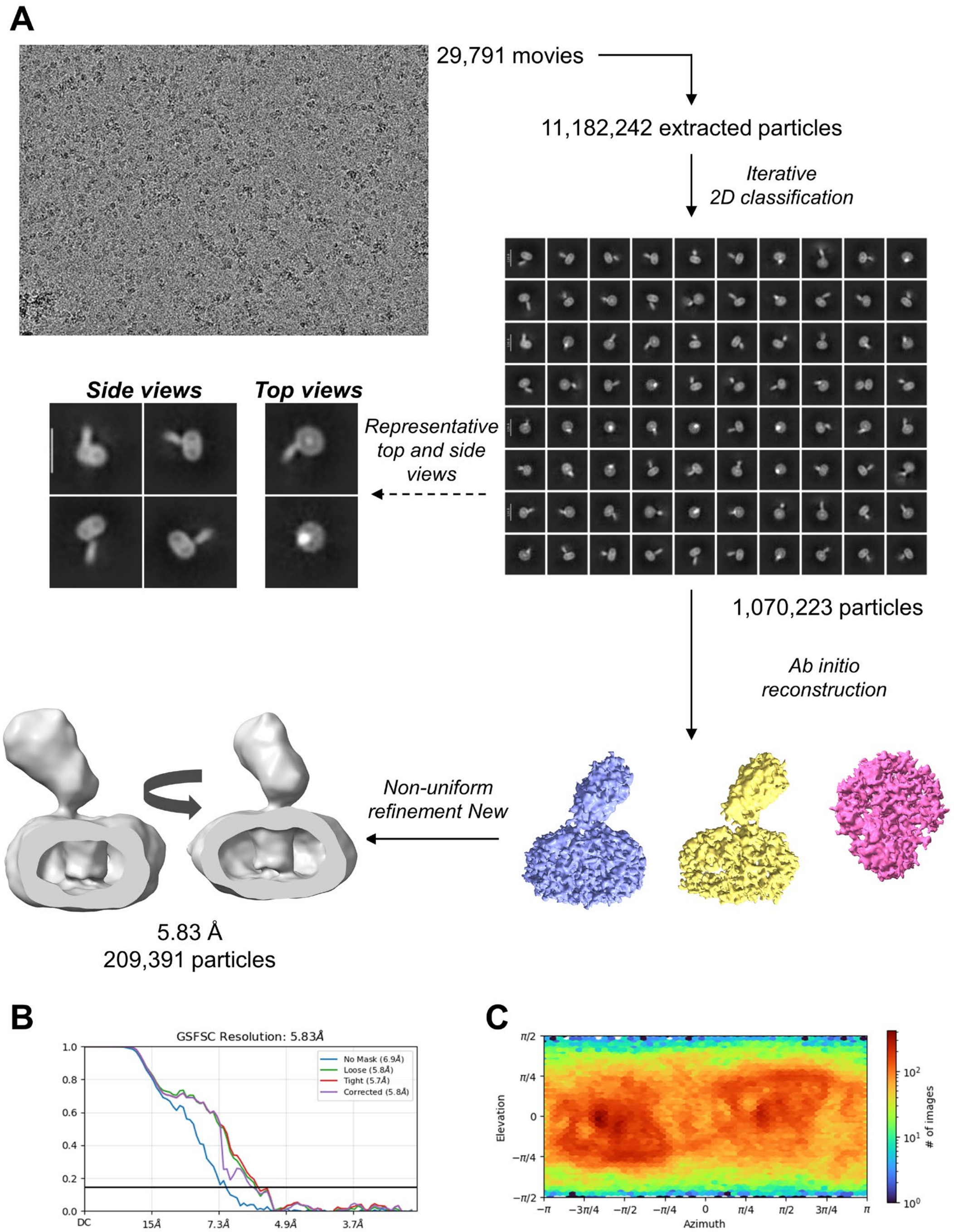

**Supplementary Figure 9. (A)** Representative movie and cryo-EM workflow for data processing. Details are described in the Methods. The dotted arrow highlights representative 2D class averages from top and side views highlighting the positioning of the toxin on one edge of the receptor**. (B)** Gold-standard Fourier shell correlation (GSFSC) curve, and **(C)** particle angular distribution for the non-uniform refinement map of the ACKR1-HlgA complex.

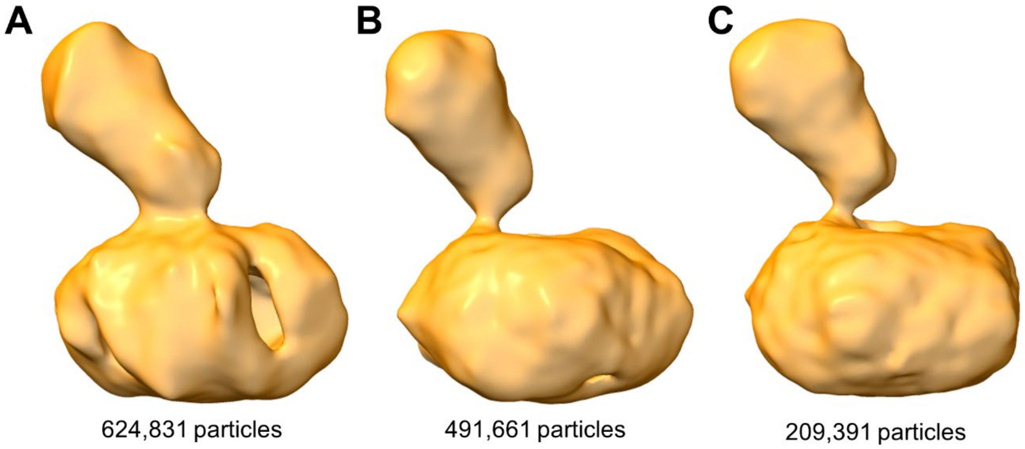

**Supplementary Figure 10. (A-C)** Non-uniform cryo-EM densities, composed of different number of particles, reveal the global positioning of the toxin relative to the receptor. The maps reveal that the toxin binds the receptor on one side at an angle of about 65°.

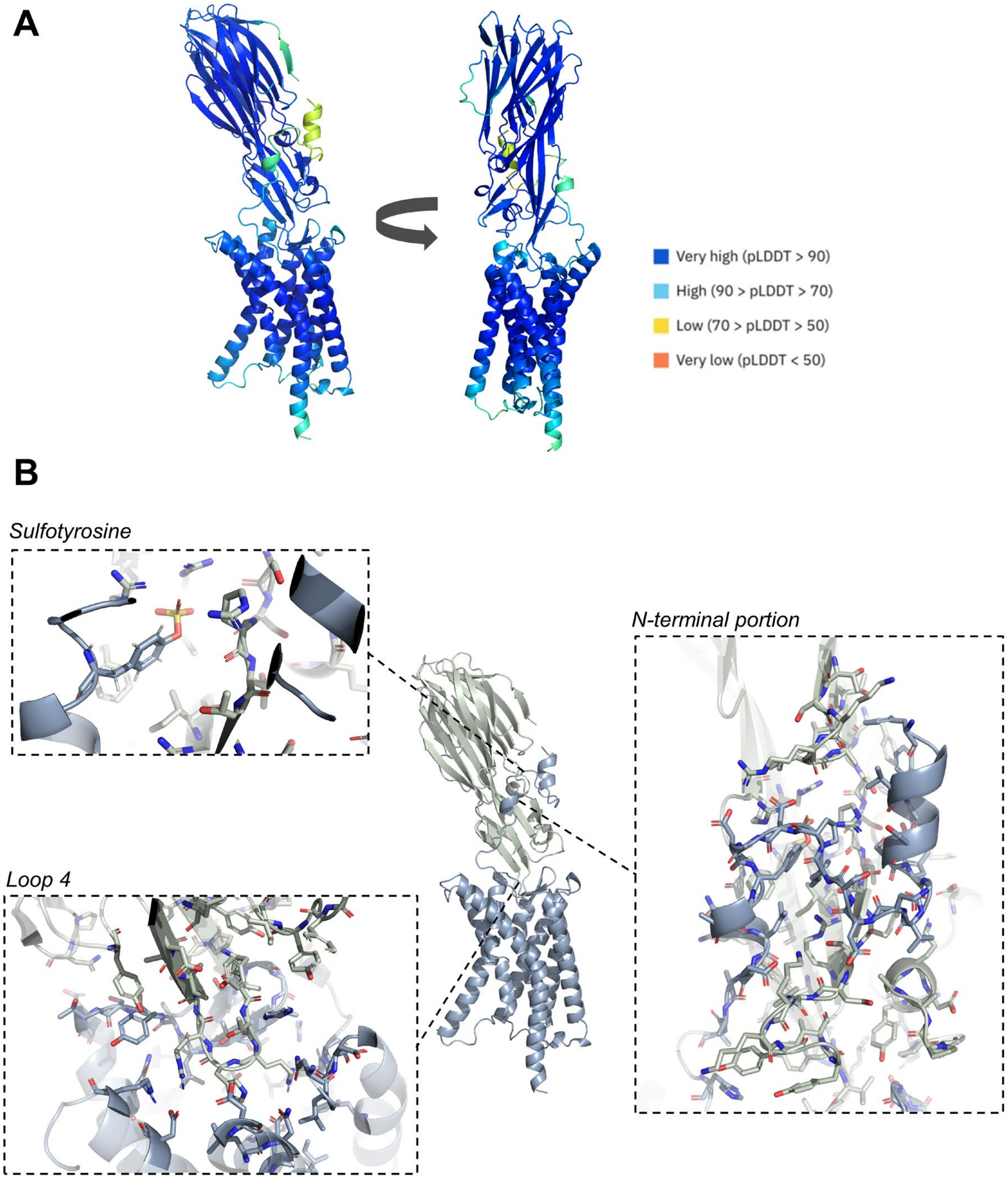

**Supplementary Figure 11. (A)** Protenix V2 model of ACKR1-HlgA complex and corresponding pLDDT score. **(B)** Interactions between ACKR1 and HlgA, with extensive contacts between the N-terminal portion of the receptor and the toxin, especially at the sulfotyrosine site of ACKR1 and the loop 4 of HlgA. All the interactions are listed in Supplementary Table 3.

### *Supplementary tables*

| Sample | Rg (nm) | Theoretical MW (kDa) | | Vc-based MW (kDa) | EOM/GAJOE χ² | | | | |
|---|---|---|---|---|---|---|---|---|---|
| | | *apo* | +4 $^{sulfo}$P | | *apo* | +1 $^{sulfo}$P | +2 $^{sulfo}$P | +3 $^{sulfo}$P | +4 $^{sulfo}$P |
| HlgA *apo* | 2.90 ± 0.02 | 36 | *n/a* | 36.6 | 1.516 | *n/a* | *n/a* | *n/a* | *n/a* |
| HlgA + $^{sulfo}$P | 2.84 ± 0.02 | 36 | 42 | 43 | *n/a* | 2.982 | 2.215 | 1.534 | 1.468 |
| HlgB *apo* | 2.63 ± 0.01 | 35.6 | *n/a* | 30.3 | 1.263 | *n/a* | *n/a* | *n/a* | *n/a* |
| HlgB + $^{sulfo}$P | 3.02 ± 0.03 | 35.6 | 41.6 | 37.6 | *n/a* | 1.75 | 1.552 | 1.341 | 1.323 |

**Supplementary Table 1.** SAXS parameters for the toxins in their *apo* form and in the presence of saturating conditions of $^{sulfo}$P. Radius of gyration (Rg) was determined by Guinier analysis. Theoretical molecular weight (MW) was calculated from the protein sequence and the MW of the $^{sulfo}$P (1519 Da). Molecular weight was also estimated from the Porod volume-corrected parameter (Vc-based MW). Ensemble Optimization Method (EOM) analysis was performed using GAJOE to assess conformational flexibility, with the reported χ² reflecting the goodness of fit between the selected ensemble and the experimental scattering data. EOM/GAJOE χ² values are shown for ensembles built with 1-4 copies of $^{sulfo}$P in complex with the respective toxin (as predicted by protenix). The best (lowest χ²) fit for each peptide-bound sample is highlighted in green.

| | ACKR1-HIgA<br>EMD-59457 |
|---|---|
| **Data collection and processing** | |
| Magnification | 105,000 |
| Voltage (kV) | 300 |
| Electron exposure (e/Å$^2$) | 39.5 |
| Defocus range (μm) | -1.0 to -2.2 |
| Pixel size (Å) | 0.84 |
| Symmetry imposed | C1 |
| **Local refinement** | |
| Particle used | 209,391 |
| Map resolution (Å) | 5.8 |
| FSC threshold | 0.143 |

**Supplementary Table 2.** Cryo-EM data collection and processing.

| HIgA | | ACKR1 | |
|---|---|---|---|
| ***Region*** | ***Residue*** | ***Residue*** | ***Region*** |
| D-region | T50, S51 | F22 | N-ter |
| | K52 | F22, V25 | |
| | R53 | D38, D40 | |
| | L54 | D38 | |
| | A55 | F22, V25, W26 | |
| | I56, T57 | F22 | |
| Divergent region/Loop1 | F82 | F22, W26 | |
| | I83 | F22 | |
| | S84 | F22, V25, W26, Y30 | |
| | S85 | Y30, Y41 | |
| | R86 | V25, W26, S28-Y30, F36-D38, Y41 | |
| | T87 | S29, Y30, F46 | |
| | T88 | S29-G31, N33, D34, F36, P37 | |
| | Y89 | S29-V32, D34 | |
| | S90 | V32, D34 | |
| | D91 | D34 | |
| | L92 | L45-A47 | |
| | K93 | D34, L45, E46 | |
| | K94 | E46 | |
| | Y95 | A48 | |
| | R100 | F36, L45 | |
| | I102 | F36, Y41, L45 | |
| Divergent region/Loop2 and Loop3 | N196 | S200-E202 | ECL2 |
| | Y205 | Y199 | |
| | D210 | A190, S191, I198 | |
| | P211 | S191, G192 | |
| | A215, A216 | V32 | N-ter |
| | R217 | Y30, V32 | |
| | Y218 | G192 | ECL2 |
| | D223, I230 | Y30 | N-ter |
| | Q231, S232 | W26, Y30 | |
| Site 1 | R260 | D38, Y41 | |
| | M262 | F36, Y41 | |
| Bottom β-pleated blade 2 | A264 | N44, L45 | |
| | Y266 | N44-A47 | |
| Divergent region/Loop4 | Y268 | A49 | |
| | V269 | S191-L194 | ECL2 |
| | T270 | L194, T196 | |
| | **R271** | N55, L56, D58, F65 | Top TM1 |
| | **R271** | L116, A117, P118 | Top TM2/ECL1 |
| | **R271** | Y133 | Pocket at TM3 |

| | | | |
|---|---|---|---|
| | **R271** | L194, C195, T196 | ECL2 |
| | **R271** | E290 | Pocket at TM7 |
| Divergent region/Loop4 | **H272** | T196, L197 | ECL2 |
| | **H272** | L286, N287, E290 | Pocket at TM7 |
| | **R273** | T196, L197, Y198 | ECL2 |
| | **R273** | Q207 | Pocket at TM5 |
| | **R273** | D263, R267 | Pocket at TM6 |
| | **R273** | L286 | Pocket at TM7 |
| | L274 | S191, T196-Y199 | ECL2 |
| | D277 | A49 | N-ter |
| Bottom β-pleated blade 2 | K279 | N44, A47 | |
| | A282 | N44 | |
| Site 1 | F283 | D40, Y41, N44, L45 | |
| | R286 | D38, D40, Y41 | |

**Supplementary Table 3.** Interface contacts between HlgA and ACKR1 in the generated model. Residue-Residue contacts were identified using a distance cutoff of 4 Å between Cα atoms.

| | | **Sequence** |
|---|---|---|
| ***Peptide*** | | |
| $^{sulfo}$P | | Ac-DSFPDGD$^{sulfo}$YGANLE-NH2 |
| scr$^{sulfo}$P | | Ac-S$^{sulfo}$YLDPDGFNDGAE-NH2 |
| P | | Ac-DSFPDGDYGANLE-NH2 |
| ***Protein*** | ***Plasmid*** | |
| HlgA-(His)8-TST | pET41(a)+ | MGS$^{30}$ENKIEDIGQGAEIIKRTQDITSKRLAITQNIQFDF<br>VKDKKYNKDALVVKMQGFISSRTTYSDLKKYPYIKR<br>MIWPFQYNISLKTKDSNVDLINYLPKNKIDSADVSQK<br>LGYNIGGNFQSAPSIGGSGSFNYSKTISYNQKNYVT<br>EVESQNSKGVKWGVKANSFVTPNGQVSAYDQYLFA<br>QDPTGPAARDYFVPDNQLPPLIQSGFNPSFITTLSHE<br>RGKGDKSEFEITYGRNMDATYAYVTRHRLAVDRKH<br>DAFKNRNVTVKYEVNWKTHEVKIKSITPK$^{309}$HHHHH<br>HHHWSHPQFEKGGGSGGGSGGGSWSHPQFEK |
| HlgB-(His)8 | pET41(a)+ | MGS$^{26}$AEGKITPVSVKKVDDKVTLYKTTATADSDKFKI<br>SQILTFNFIKDKSYDKDTLVLKATGNINSGFVKPNPN<br>DYDFSKLYWGAKYNVSISSQSNDSVNVVDYAPKNQ<br>NEEFQVQNTLGYTFGGDISISNGLSGGLNGNTAFSE<br>TINYKQESYRTTLSRNTNYKNVGWGVEAHKIMNNG<br>WGPYGRDSFHPTYGNELFLAGRQSSAYAGQNFIAQ<br>HQMPLLSRSNFNPEFLSVLSHRQDGAKKSKITVTYQ<br>REMDLYQIRWNGFYWAGANYKNFKTRTFKSTYEID<br>WENHKVKLLDTKETENNK$^{325}$LEHHHHHHHH |
| HlgA$_{\Delta site1}$-(His)8-TST | pET41(a)+ | MGS$^{30}$ENKIEDIGQGAEIIKRTQDITSKRLAITQNIQFDF<br>VKDKKYNKDALVVKMQGFISSGTTYSDLKKYPYIKR<br>MIWPFQYNISLKTKDSNVDLINYLPKNKIDSADVSQK<br>LGYNIGGNFQSAPSIGGSGSFNYSKTISYNQKNYVT<br>EVESQNSKGVKWGVKANSFVTPNGQVSAYDQYLFA<br>QDPTGPAARDYFVPDNQLPPLIQSGFNPSFITTLSHE<br>RGKGDKSEFEITYGRNMDATYAYVTRHRLAVDRKH<br>DAFKNENVTVKYEVNWKTHEVKIKSITPK$^{309}$HHHHHH<br>HHWSHPQFEKGGGSGGGSGGGSWSHPQFEK |
| δHlgA-(His)8 | pET41(a)+ | MGS$^{30}$ENKIEDIGQGAEIIKRTQDITSKRLAITQNIQFDF<br>VKDKKYNKDALVVKMQGFISSRTTYSDLKKYPYIKR<br>MIWPFQYNISLKTKDSNVDLINYLPKNKIDSADVSQK<br>LNFQSAPSIGGSGSFNYSKTISYNQKNYVTEVESQN<br>SKGVKWGVKANSFVTPNGQVSAYDQYLFAQDPTG<br>PAARDYFVPDNQLPPLIQSGFNPSFITTLSHERGKG<br>DKSEFEITYGRNMDATYAYVTRHRLAVDRKHDAFKN<br>RNVTVKYEVNWKTHEVKIKSITPK$^{303}$LEHHHHHHHH |

| | | |
|---|---|---|
| δHIgB-(His)8 | pET41(a)+ | MGS$^{26}$AEGKITPVSVKKVDDKVTLYKTTATADSDKFKI<br>SQILTFNFIKDKSYDKDTLVLKATGNINSGFVKPNPN<br>DYDFSKLYWGAKYNVSISSQSNDSVNVVDYAPKNQ<br>NEEFQVQNTLGYTFGGDISISNGLSNTAFSETINYKQ<br>ESYRTTLSRNTNYKNVGWGVEAHKIMNNGWGPYG<br>RDSFHPTYGNELFLAGRQSSAYAGQNFIAQHQMPL<br>LSRSNFNPEFLSVLSHRQDGAKKSKITVTYQREMDL<br>YQIRWNGFYWAGANYKNFKTRTFKSTYEIDWENHK<br>VKLLDTKETENNK$^{320}$LEHHHHHHHH |
| Flag-ACKR1 | pFastBac | DYKDDDDKASSGYVLQ$^{8}$AELSPSTENSSQLDFEDV<br>WNSSYGVNDSFPDGDYGANLEAAAPCHSCNLLDDS<br>ALPFFILTSVLGILASSTVLFMLFRPLFRWQLCPGWP<br>VLAQLAVGSALFSIVVPVLAPGLGSTRSSALCSLGYC<br>VWYGSAFAQALLLGCHASLGHRLGAGQVPGLTLGL<br>TVGIWGVAALLTLPVTLASGASGGLCTLIYSTELKAL<br>QATHTVACLAIFVLLPLGLFGAKGLKKALGMGPGPW<br>MNILWAWFIFWWPHGVVLGLDFLVRSKLLLLSTCLA<br>QQALDLLLNLAEALAILHCVATPLLLALFCHQATRTLL<br>PSLPLPEGWSSHLDTLGSKS$^{336}$LPV |
| Flag-SNAP-ACKR1 | pcDNA 3.1 | DYKDDDDASGDKDCEMKRTTLDSPLGKLELSGCEQ<br>GLHEIKLLGKGTSAADAVEVPAPAAVLGGPEPLMQA<br>TAWLNAYFHQPEAIEEFPVPALHHPVFQQESFTRQV<br>LWKLLKVVKFGEVISYQQLAALAGNPAATAAVKTALS<br>GNPVPILIPCHRVVSSSGAVGGYEGGLAVKEWLLAH<br>EGHRLGKPGLARGIRASSGYVLQ$^{8}$AELSPSTENSSQ<br>LDFEDVWNSSYGVNDSFPDGDYGANLEAAAPCHSC<br>NLLDDSALPFFILTSVLGILASSTVLFMLFRPLFRWQL<br>CPGWPVLAQLAVGSALFSIVVPVLAPGLGSTRSSAL<br>CSLGYCVWYGSAFAQALLLGCHASLGHRLGAGQVP<br>GLTLGLTVGIWGVAALLTLPVTLASGASGGLCTLIYS<br>TELKALQATHTVACLAIFVLLPLGLFGAKGLKKALGM<br>GPGPWMNILWAWFIFWWPHGVVLGLDFLVRSKLLL<br>LSTCLAQQALDLLLNLAEALAILHCVATPLLLALFCHQ<br>ATRTLLPSLPLPEGWSSHLDTLGSKS$^{336}$LPV |
| Flag-SNAP-ACKR1$_{\Delta Nter}$ | pcDNA 3.1 | DYKDDDDASGDKDCEMKRTTLDSPLGKLELSGCEQ<br>GLHEIKLLGKGTSAADAVEVPAPAAVLGGPEPLMQA<br>TAWLNAYFHQPEAIEEFPVPALHHPVFQQESFTRQV<br>LWKLLKVVKFGEVISYQQLAALAGNPAATAAVKTALS<br>GNPVPILIPCHRVVSSSGAVGGYEGGLAVKEWLLAH<br>EGHRLGKPGLARGIRASSGYVLQ$^{8}$AELSPSTENSSQ<br>LDFEDVWNSSYGVNAAAPCHSCNLLDDSALPFFILT<br>SVLGILASSTVLFMLFRPLFRWQLCPGWPVLAQLAV<br>GSALFSIVVPVLAPGLGSTRSSALCSLGYCVWYGSA<br>FAQALLLGCHASLGHRLGAGQVPGLTLGLTVGIWGV |

| | | |
|---|---|---|
| | | AALLTLPVTLASGASGGLCTLIYSTELKALQATHTVA<br>CLAIFVLLPLGLFGAKGLKKALGMGPGPWMNILWAW<br>FIFWWPHGVVLGLDFLVRSKLLLLSTCLAQQALDLLL<br>NLAEALAILHCVATPLLLALFCHQATRTLLPSLPLPEG<br>WSSHLDTLGSKS$^{323}$LPV |
| Flag-SNAP-μOR | pcDNA 3.1 | DYKDDDDASGDKDCEMKRTTLDSPLGKLELSGCEQ<br>GLHEIKLLGKGTSAADAVEVPAPAAVLGGPEPLMQA<br>TAWLNAYFHQPEAIEEFPVPALHHPVFQQESFTRQV<br>LWKLLKVVKFGEVISYQQLAALAGNPAATAAVKTALS<br>GNPVPILIPCHRVVSSSGAVGGYEGGLAVKEWLLAH<br>EGHRLGKPGLGAS$^{1}$MDSSAGPGNISDCSDPLAPASC<br>SPAPGSWLNLSHVDGNQSDPCGPNRTGLGGSHSL<br>CPQTGSPSMVTAITIMALYSIVCVVGLFGNFLVMYVI<br>VRYTKMKTATNIYIFNLALADALATSTLPFQSVNYLM<br>GTWPFGNILCKIVISIDYYNMFTSIFTLCTMSVDRYIA<br>VCHPVKALDFRTPRNAKIVNVCNWILSSAIGLPVMFM<br>ATTKYRQGSIDCTLTFSHPTWYWENLLKICVFIFAFIM<br>PVLIITVCYGLMILRLKSVRMLSGSKEKDRNLRRITRM<br>VLVVVAVFIVCWTPIHIYVIIKALITIPETTFQTVSWHFC<br>IALGYTNSCLNPVLYAFLDENFKRCFREFCIPTSSTIE<br>QQNSARIRQNTREHPSTANTVDRTNHQLENLEAETA<br>PLP$^{398}$ |

**Supplementary Table 4.** Peptide and protein sequences used in this study.
